# The Ephrin receptor A2 and Roundabout Guidance Receptor 1 heterodimer: A potential theranostic for squamous cell carcinomas

**DOI:** 10.1101/2020.04.09.034405

**Authors:** Ka M. Pang, Saumya Srivastava, Mari Iida, Michael Nelson, Jiayi Liu, Arin Nam, Jiale Wang, Isa Mambetsariev, Atish Mohanty, Nellie McDaniel, Amita Behal, Prakash Kulkarni, Deric L. Wheeler, Ravi Salgia

**Affiliations:** Department of Surgery, City of Hope National Medical Center, Duarte, CA, USA; Department of Medical Oncology & Therapeutics Research, City of Hope National Medical Center, Duarte, CA, USA; Department of Human Oncology, University of Wisconsin School of Medicine and Public Health, Madison, WI, USA; Light Microscopy Core, City of Hope National Medical Center, Duarte, CA, USA

## Abstract

Squamous cell carcinomas (SCC) of the lung (LSCC) and head and neck (HNSCC) are very prevalent with poor prognosis and limited treatment options. In both cancer types, Ephrin receptor A2 (EPHA2) is known to be overexpressed and exhibit opposing effects via two distinct signaling mechanisms. While it can inhibit cancer cell survival and migration by ligand-dependent signaling through tyrosine kinase phosphorylation at Y588 and Y772, it can promote tumor progression and cell migration in a ligand-independent manner via phosphorylation at S897. Variable ABnormal morphology (VAB-1) is the *C. elegans* ortholog of the human ephrin receptor (EPHR) that interacts genetically and biochemically in a dose-dependent manner with the axon guidance receptor, SAX3, the worm ortholog of ROBO. Double mutants of *vab-1*(EPHR)/*sαx-3*(ROBO) are synthetic lethal, underscoring the interaction between the two signaling pathways which prompted us to investigate their role in SCC. Using biochemical and biophysical techniques, we show that EPHA2 and ROBO1 reside in the same complex and interact physically to form a functional heterodimer in LSCC and HNSCC. Furthermore, we show that treating squamous cells with the SLIT2, ligand of ROBO1, hinders phosphorylation of EPHA2 at S897, and thereby, attenuates cell proliferation. Interestingly, SLIT2 can interact with EPHA2 and attenuate the proliferation of cells that have low ROBO1 expression. Additionally, SLIT2 can act synergistically with the EPHA2 inhibitor, Ensartinib to attenuate cell growth in LSCC and HNSCC cells. Taken together, the data suggest that SLIT2 may serve as a novel therapeutic for LSCC and HNSCC. Here, we propose to stratify patients for treatment with SLIT2 and/or Ensartinib, based on their EPHA2 and ROBO1 expression levels in the diseased tissue. Thus 85% of LSCC cases can be treated with combination of SLIT2+Ensartinib and 55% of HNSCC cases can be treated with either SLIT2 or Ensartinib. Furthermore, EPHA2 and ROBO1 may represent novel theranostics in these two diseases.

**One sentence summary:** Heterodimerization of EPHA2 and ROBO1 receptors attenuates growth of squamous cell carcinomas of the lung and head and neck.

## Introduction

Lung cancer is the leading cause of cancer-related deaths worldwide with 1.76 million deaths in 2018 (*1*). With an estimated 1.8 million new lung cancer cases, it accounts for about 13% of total cancer diagnoses (*2*). Non-small cell lung cancer (NSCLC) represents around 85% of all lung cancers with the majority of patients presenting with advanced stages of the disease when diagnosed. Squamous cell carcinoma (SCC) is the second most common histology in NSCLC accounting for 20–30% of all NSCLC cases and more often than not, presents with advanced stage disease at diagnosis (*3*). Head and neck cancers represent the sixth most common cancer worldwide and accounts for approximately 650,000 new cases and more than 350,000 deaths every year. In the United States, head and neck cancer accounts for 3 percent of all cancers, with estimated 53,000 Americans developing head and neck cancer annually and 10,800 dying from the disease. Approximately 90% of head and neck cancers are squamous cell carcinomas (HNSCC) that arise from the mucosal surfaces of the oral cavity, oropharynx and larynx. Tobacco use, alcohol consumption, Epstein-Barr virus infection and HPV infection are some of the risk factors that are known to contribute to the development of almost 80% of the HNSCC cases diagnosed globally (*4,5*). However, there are limited treatment options for advanced SCC, both in first line and relapsed/refractory settings. In the last few years, several new drugs that include an anti-EGFR monoclonal antibody (necitumumab) in combination with standard chemotherapy, and immune-checkpoint inhibitors such as nivolumab, pembrolizumab, or atezolizumab have been approved for treating SCC. Although the initial response to these drugs is encouraging, the patients eventually acquire resistance during the course of treatment, emphasizing the need for new drug targets and more efficacious drugs (*6*).

The ephrin receptors (EPHR) and their ligands, the ephrins, play critical roles in a broad range of biological processes including neuronal pathfinding, angiogenesis, T cell activation, and stem cell maintenance (*7,8*). At the cellular level, the EPHR pathway regulates polarity, motility, proliferation and survival in normal as well as neoplastic conditions (*9,10*). In humans, ephrins and ephrin receptors constitute the largest receptor tyrosine kinase family with a total of 14 members. Upregulation of EPH receptors, particularly EPHA2 and EPHB4, has been broadly implicated in the growth and metastasis of solid tumors, including NSCLC (*11–17*). High levels of EPHA2 have correlated with brain metastasis, disease relapse, and overall poor patient survival in lung cancers (*18*). In addition, EPHA2 is involved in acquired resistance to EGFR tyrosine kinase inhibitors (TKIs) and anti-EGFR antibodies (*19–21*), further highlighting the role of EPHA2 in tumor progression. Ephrins, are classified into either class A or B depending upon how they bind to the cell membrane. Class A ephrins (ephrin A1 to A5) are tethered to the cell membrane via a Glycosyl-phosphatidylinositol (GPI) anchor, while class B ephrins (ephrin B1 to B3) are integral membrane proteins that contain a single transmembrane domain. EPH receptors are classified according to the class of ligand they bind to. There are 8 class A receptors (EPHA1 to A8) and 6 class B receptors (EPHB1 to B6) (*15*). Both the ligands and the receptors are membrane bound and typically elicit signaling through cell-cell contact that leads to clustering of the ligand-receptor complexes. In addition, the binding of the ligand to the receptor triggers downstream events in the ligand-bearing cells, referred to as reverse signaling (*22,15–16*). Among the EPHRs, EPHA2 has been shown to possess both ligand-dependent as well as ligand-independent activities (*23,24*). While ligand-mediated activation of EPHA2 was shown to inhibit cell motility, the ligand independent activities enhance invasiveness and viability, and are linked to tumor progression (*23, 25–27*). Importantly, S897 phosphorylation by AKT or RSK is a key mediator of the ligand-independent activities (*23,28*) leading to tumorigenesis. In *C. elegans,* the ortholog of the human ephrin receptor, VAB-1, and its ligand, ephrin, EFN-1, function as cues in neuroblasts for proper neuronal pathfinding and in epidermal movements during embryogenesis. Mutants of the VAB-1 and ephrin exhibit variable degree of defects in embryonic arrest, larval arrest and morphogenesis with high percentage of animal showing notch head phenotype (*29, 30*). The Ephrin/VAB-1 pathway interacts genetically with the conserved axon guidance receptor SAX-3/ROBO pathway in a dose-dependent fashion and showed synthetic embryonic lethal phenotype (*31,32*). SAX-3/ROBO null mutants display embryonic defects independent of VAB-1/EPHR mutants. Interestingly double mutants of SLT-1, a ligand of SAX-3 (ROBO) and VAB-1 also show enhanced cell migration defects in comparison with VAB-1/EPHR or SLT-1 single mutants indicating cross talk between the two pathways. In addition, the tyrosine kinase domain of VAB-1 physically binds to the intracellular domain of SAX-3, specifically at the juxtamembrane and the conserved cytoplasmic region 1 (CC1) (*31*).

Roundabout guidance receptors (ROBO) and their ligands, SLITs, were first identified as important signaling molecules involved in the neuronal development in *Drosophila* (*33,34*). Since then, this function in neuronal development has been found to be highly conserved in metazoans. In addition, new functions of the SLIT-ROBO pathway have been discovered in angiogenesis, and in the development of the lung, mammary glands and kidneys. (*35–40*). Recent studies have also implicated the SLIT-ROBO pathway in cancer progression and metastasis (*42–44*). There are four ROBOs (ROBO 1-4) and three SLITs (SLIT 1-3) in mammals that can bind to different ROBO receptors with differing affinities. All ROBO receptors contain a single transmembrane domain with several weakly conserved cytoplasmic (CC) domains and no clear functionally defined domain in the cytoplasmic tail. Therefore, additional signaling molecules are probably involved in directing cellular activities (*45,43,40*). ROBO1 overexpression in lung cancer has been correlated with better patient outcome (**Table 1 and Supplementary Figure 2A**). Suppression of SLIT2 was associated with advanced pathological stage and a poor survival rate among lung cancer patients (*43*). Despite the correlation between expression levels of EPHA2, ROBO1 and SLIT2 and tumorigenesis and clinical outcome in lung cancer patients, the translational potential of this clinical research has not been fully explored.

**Table 1.** Survival plot of NSCLC cases with ROBO1 high expression versus ROBO1 low expression

|  | # of cases | Medium Months survival |
| --- | --- | --- |
| **ROBO1 high expression** | 55 | 129.08 |
| **ROBO1 low expression** | 54 | 31.64 |

**Table 2.** Survival plot of NSCLC cases with alteration in ROBO1 gene versus no ROBO1 gene alterations

|  | # of cases,<br>Total | # of cases,<br>Deceased | Medium Months survival |
| --- | --- | --- | --- |
| **Cases with Alternation in ROBO1** | 14 | 7 | 8.44 |
| **Cases without Alternation in ROBO1** | 189 | 56 | 45.3 |

Here, we have investigated the roles of EPHA2 and ROBO1 in SCCs of the lung and head and neck. Our results demonstrate that EPHA2 can physically interact by heterodimerizing with ROBO1 and this interaction is stabilized in the presence of SLIT2 which in turn attenuates cellular proliferation. Thus, SLIT2 is a potential novel therapeutic for squamous cell carcinomas of the lung and head and neck, and EPHA2 and ROBO1 may represent novel theranostics in these two diseases.

## Results

### ROBO1 and EPHA2 are not synthetic lethal in LSCC and HNSCC cell lines

In *C. elegans,* double mutants of *sax-3* in the SLIT-ROBO and *vab-1* the Ephrin-EPH pathway showed synthetic lethal phenotype in embryonic stage suggesting that there is crosstalk between the two pathways (*31*). This interaction presents an attractive window of opportunity for developing targeted therapy against EPH-ROBO pathway, if the underlying mechanism is conserved in SCC. Therefore, we first determined whether a knockdown or pharmacological inhibition of the EPH receptor (*vab-1*) in *C. elegans* exhibited a synthetic lethal phenotype with ROBO mutant (*sax-3*). Consistent with the phenotype of genetic double mutants, knocking down *vab-1* in *sax-3* or knocking *sax-3* in *vab-1* mutants exhibited the synthetic lethal phenotype (**Supplementary Fig. 1A and B**). Moreover, treating *sax-3* worms with ALW-II-41-27, a small molecule Eph family tyrosine kinase inhibitor, also enhanced lethality (**Supplementary Fig. 1C**), indicating that ALW-II-41-27 inhibited the ephrin receptor of *C. elegans* (*46, 18, 47*).

Next, to determine if EPHA2 is important for the survival of lung squamous cell and adenocarcinoma carcinoma cells, we inhibited EPHA2 by shRNA knockdown as well as ALW-II-41-27 treatment in human cell lines (*46, 18, 47*). We knocked down EPHA2 using shRNA in three squamous cell carcinoma (SK-MES-1, H2170, SW900) and two adenocarcinomas (A549, SK-LU-1) cell lines as well as control normal lung epithelial BEAS-2B cells and measured total viability at 72 h post-transfection. All five cell lines showed varying degrees of inhibition, while control BEAS-2B cells showed minimal growth inhibition (**Fig. 1A**). A549, SK-LU-1 and SK-MES-1 showed 65%, 38% and 52% inhibition respectively whereas H2170 and SW900 showed only 20% inhibition. Similarly, treatment of cells with ALW-II-27-41 resulted in stronger inhibition of NSCLC cells (IC50 range from 134nM to 1253nM) relative to control BEAS-2B cells (IC50=1533nM) (**Fig. 1B**). These results support the idea that EPHA2 is involved in positive signaling for NSCLC cell proliferation.

**Figure 1.**
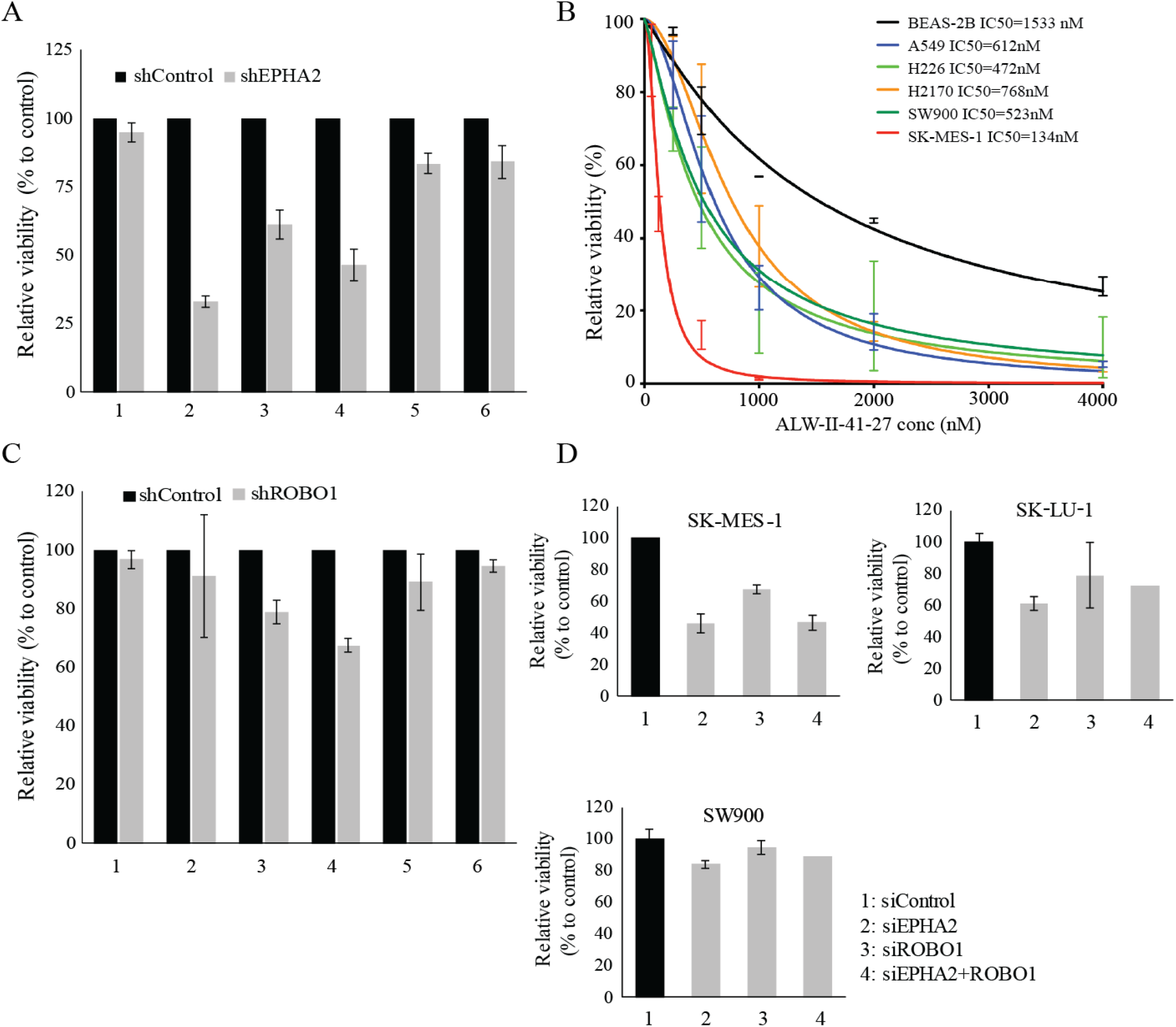
shRNA mediated knock down of EPHA2 and ROBO1 to study effect on cellular proliferation in lung cancer cell lines. (A): Knock down of EPHA2 via shRNA has variable effects on cellular proliferation among different cell lines in comparison to control cells BEAS-2B. (B): Chemical ablation of EPHA2 using ALW-II-41-27(EPH inhibitor) shows significant inhibition in proliferation in all lung cancer cell lines in comparison to control lung epithelial cells BEAS2B indicated by values of IC50.(C): Knock down of ROBO1 via shRNA have variable effects on cellular proliferation. With most cell lines not significantly affected. (D): EPHA2 and ROBO1 shRNA double knockdown in NSCLC cancer cell lines is not synergistic. 1: BEAS-2B; 2: A549; 3: SK-LU-1; 4: SK-MES-1; 5: H2170; 6: SW900.

To further investigate the functional relationship between ROBO1 and EPHA2 in NSCLC, we first determined whether ROBO1 knockdown also affects cell proliferation in various NSCLC cell lines. Knocking down ROBO1 showed only moderate inhibition in SK-LU-1 (21%) and SK-MES-1 (33%) cells while no significant inhibition was observed in A549, H2170 and SW900 cells (**Fig. 1C**). We then asked if knocking down both EPHA2 and ROBO1 would show an enhancement of the inhibition shown in SK-MES-1, SK-LU-1 and SW900. In all three cell lines, no additive effect was observed (**Fig. 1D**). In contrast, a small attenuation of the inhibition caused by shEPHA2 was observed when ROBO1 was also knocked down. Next a variety of, head and neck cancer cell lines and cetuximab-resistant cell lines (HN30 cells) expressing high EPHA2 were investigated (**Fig. 2A, D**). EPHA2 was knocked down by siRNA and was confirmed by western blotting (**Fig. 2B, D**). However, EPHA2 knockdown had no significant effect on cell proliferation in HNSCC cell lines (except for 93-vu-147T) as determined by CCK-8 assay (**Fig. 2C, E**). Proliferation of HN-30 cell line was also not affected by siRNA knockdown of EPHA2. These observations suggest that EPHA2 and ROBO1 pathways in squamous cells may interact but may not lead to synthetic lethality as shown in *C. elegans* model.

**Figure 2.**
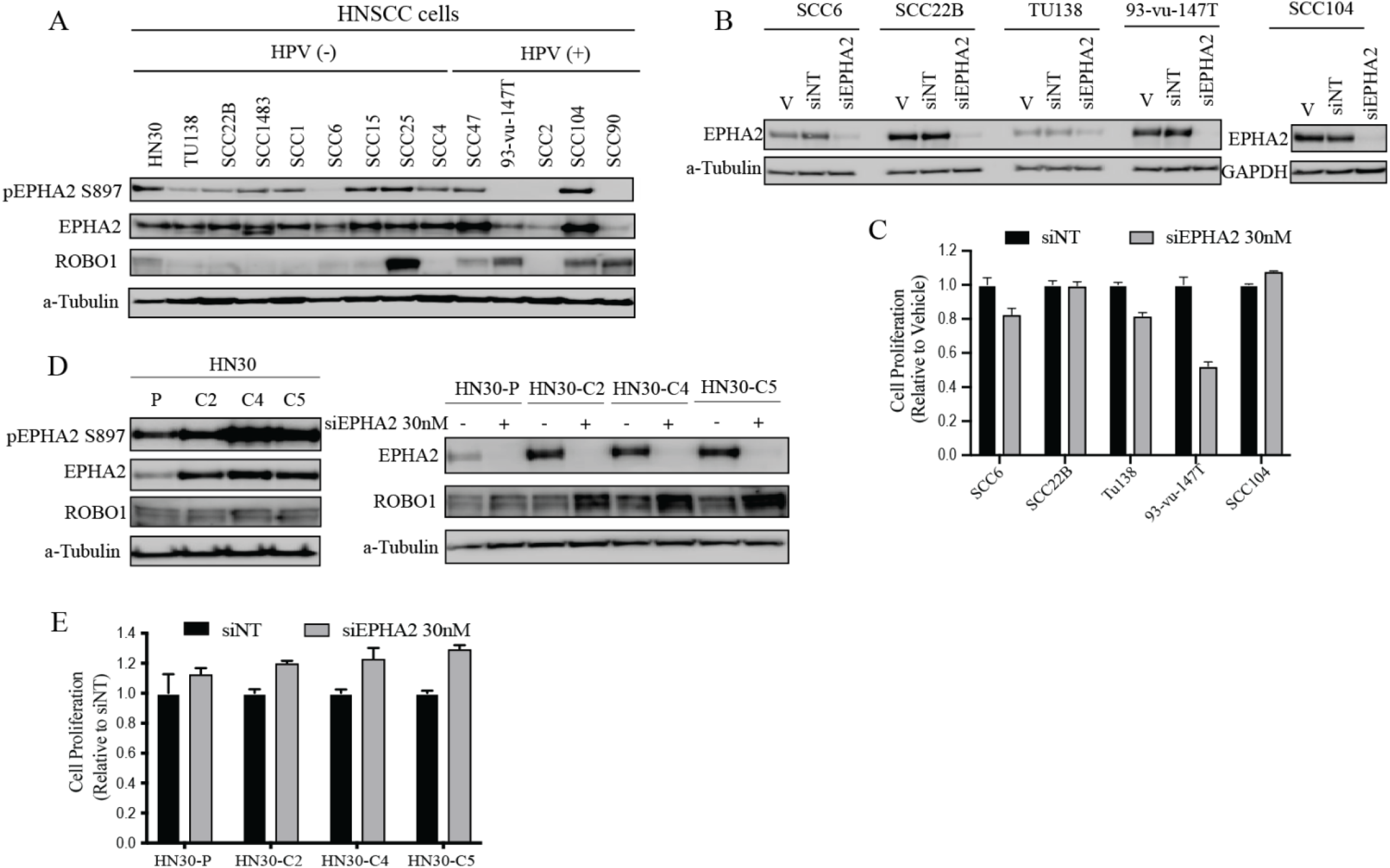
EPHA2 and ROBO1 levels in HNSCC cell lines and EPHA2 siRNA mediated knock down to study effect on cellular proliferation. (A): Levels of total EPHA2, ROBO1 and Phospho-EPHA2 S897 in head and neck cancer cell lines. (B): Levels of EPHA2 confirming knock down using siRNA. (V: Vehicle; siNT: Negative control; siEPHA2: EPHA2 knockdown). (C): CCK8 assay to determine cellular proliferation inhibition after knocking down EPHA2. (D): Levels of total EPHA2, ROBO1 and P-S897 in cetuximab resistant cells and levels of EPHA2 after knocking down EPHA2 via siRNA. (E). Relative cell number after CCK8 assay to determine cellular proliferation inhibition in EPHA2 knockdown cetuximab resistant cells.

### Expression of EPHA2 and ROBO1 receptors in LSCC and HNSCC TMAs

Our previous studies with 105 NSCLC patient samples showed that EPHA2 was overexpressed in these patients (*13*). Among the three types of NSCLCs, lung squamous cell carcinoma (LSCC) samples showed the highest expression levels of EPHA2 (n=24). To further correlate this trend, we examined EPHA2 expression using tissue microarrays (TMA), one with 64 LSCC samples and another with 80 HNSCC samples (**Fig. 3A, B**). In LSCC, 85% of the samples showed significant expression of EPHA2, while very few (~5%) showed high expression levels of only ROBO1.A quantitative evaluation of the positive staining in LSCC showed that ROBO1 and EPHA2 showed an inverse relationship where most samples with high EPHA2 expression had low ROBO1 expression (**Fig. 3A**). This correlates with the fact that higher ROBO1 levels are associated with better survival in lung cancer patients since only a relatively small percentage of lung cancer patients have good survival prognosis (**Table 1 and Supplementary Fig. 2A**). In HNSCC 55% samples show high EPHA2 and low ROBO1 expression levels (**Fig. 3B**).Adrenal gland pheochromocytoma core was used as a control core (separate from the grid at the bottom of TMA). The distribution of staining of EPHA2 and ROBO1 in the two TMAs was quantified using positive staining percent area per core using a violin plot (**Fig. 3C**). Patient stratification in LSCC and HNSCC based on TMA is summarized in (**Fig. 3D**)

**Figure 3.**
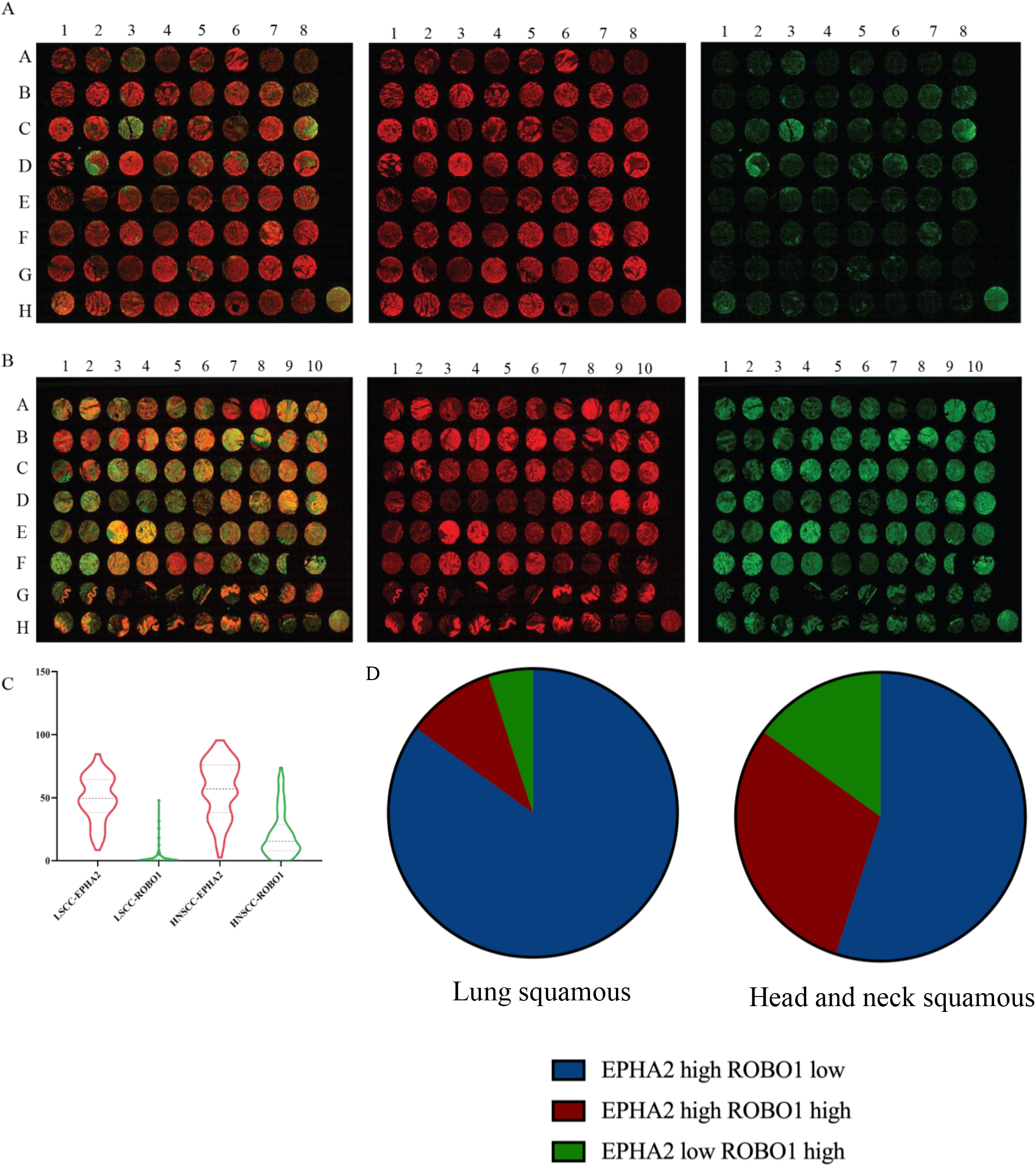
Expression pattern of EPHA2 and ROBO1 in patient tissue micro arrays in LSCC and HNSCC. (A): 64 tissue micro array panels of lung squamous cell carcinoma patient samples-TMA-LC642a Biomax. (pseudo colors show anti-EPHA2(red) and anti-ROBO1(green). The pseudo color was created by FIJI software. (B): 80 tissue micro array panels of head and neck squamous cell carcinoma patient samples-TMA_HN801B Biomax (pseudo colors show anti-EPHA2(red) and anti-ROBO1(green). (C): Quantification (described in methods) of EPHA2+ and ROBO1+ percent area, per core in overall LSCC and HNSCC TMA. Adrenal gland Pheochromocytoma tissue control core. (D): Patient stratification on the basis of EPHA2 and ROBO1 expression levels in lung squamous and head and neck squamous cancer patient tissue micro arrays.

### ROBO1/SLIT2 are present in the same complex and physically interact with EPHA2

To further probe the EPHA2/ROBO1interaction, we used fluorescent proteins to tag ROBO1 and EPHA2 and tracked their localization using confocal microscopy. ROBO1-mCherry and EPHA2-mGFP were localized to the cell surface as well as cytoplasmic vesicles (**Fig. 4A**). The merged images showed that most of the EPHA2-mGFP and the ROBO1-mCherry co-localized at the cell membrane and in cytoplasmic vesicular structures. However, some discrete loci showed either EPHA2 or ROBO1 signal (**Fig. 4A**). Z-section images revealed that both proteins localized in some discrete loci and in long extensions attached to surfaces (**Fig. 4A & Supplementary Movie 1**). In addition, we also investigated the interaction between EPHA2 (Alexa Flour 547-Red) and SLIT2 (Alexa Flour 488-Green) using confocal microscopy. We observed that EPHA2 and SLIT2 co-localized at the cell boundary and around nuclear membrane (**Fig. 4B**). To corroborate these results, we used Fluorescence Resonance Energy Transfer (FRET) and confocal microscopy and determined the proximity of ROBO1 and EPHA2. For this purpose, we tagged ROBO1 with mClover3 (ROBO1-mClover3) as the FRET donor and - EPHA2-mRuby3 (EPHA2-mRuby3) as the FRET acceptor. EPHA2-mClover3 and EPHA2-mRuby3 was used as a positive control, FRET was detected as expected with a N-FRET value of 0.347+/− 0.061. FRET was also detected between ROBO1-mCLover3 and EPHA2-mRuby3 with N-FRET value of 0.234 +/0.065 (N=23) (**Fig. 4C**). Taken together, these data provided good evidence that EPHA2 and ROBO1 are in proximity (Foster radius of ~6 nanometers) to potentially make direct contact.

**Figure 4.**
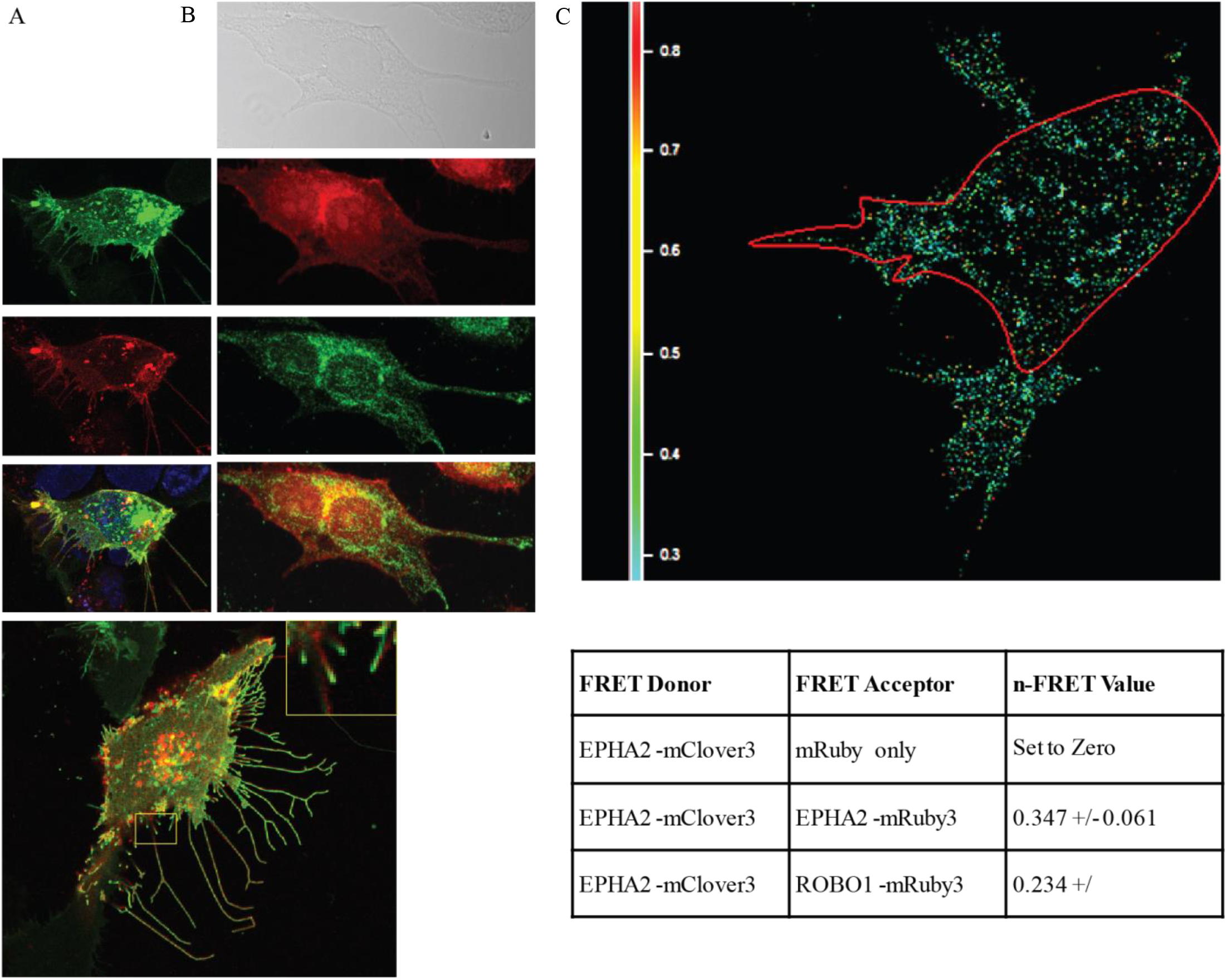
EPHA2-ROBO1 and EPHA2-SLIT2 interaction in LSCC cell line (A): EPHA2-GFP fusion(green) and ROBO1-m-cherry fusion (red) were co-expressed in cells for assessing interaction of EPHA2 and ROBO1 using confocal microscopy. The insert shows the colocalization in trailing projections. (also see supplementary video 1) (B): Coimmunofluorescence of EPHA2(red Alexa Flour 547) and SLIT2(green Alexa Flour 488) in SW900 lung squamous carcinoma cell line imaged under a confocal microscope. Yellow represents the area of co-expression of EPHA2 and SLIT2. (C): FRET analysis: EPHA2-mClover3 was used as FRET donor and EPHA2-mRuby3, ROBO1-mRuby3 both were used as the FRET acceptors. EPHA2-mClover3 and EPHA2-mRuby3 was used as a positive control. FRET values are indicated in the figure.

As additional evidence supporting interaction between EPHA2 and ROBO1, we performed coimmunoprecipitation (co-IP) experiments. Plasmids constructs with FLAG tagged ROBO-1 (ROBO1-FLAG) and HA tagged EPHA2 (EPHA2-HA) were co-transfected into HEK293 cells which express low endogenous levels of both EPHA2 and ROBO1. Indeed, in these experiments, EPHA2 protein was pulled down with ROBO1-FLAG tag IP (**Fig. 5A**). To test if the kinase activity of EPHA2 is necessary for the interaction, we generated a kinase-dead (K645R) EPHA2-KD-HA construct (*24*). Interestingly, a higher level of the kinase-dead mutant co-immunoprecipitated with ROBO1 than the wild type EPHA2, even though the kinase-dead protein expressed at a lower level than the wild type (**Fig. 5A**) suggesting that the kinase activity of EPHA2 is not required for interaction between EPHA2 and ROBO1. Similarly, when we performed the corollary experiment and used HA tag IP for HA tagged-EPHA2, a higher level of WT ROBO1 was co-immunoprecipitated with the kinase dead EPHA2 than the wild type EPHA2 (**Fig. 5B**). Additionally, ROBO1 point mutants at Y932F, a ubiquitination site, and Y1073F an ABL phosphorylation site, did not bind to WT-EPHA2. These observations suggest that ROBO1 and EPHA2 reside in the same complex and that EPHA2-ROBO1 interaction did not require the kinase activity of EPHA2. However, phosphorylation of ROBO1 is important for this interaction since phosphorylation mutants of ROBO1 are unable to interact with EPHA2.

**Figure 5.**
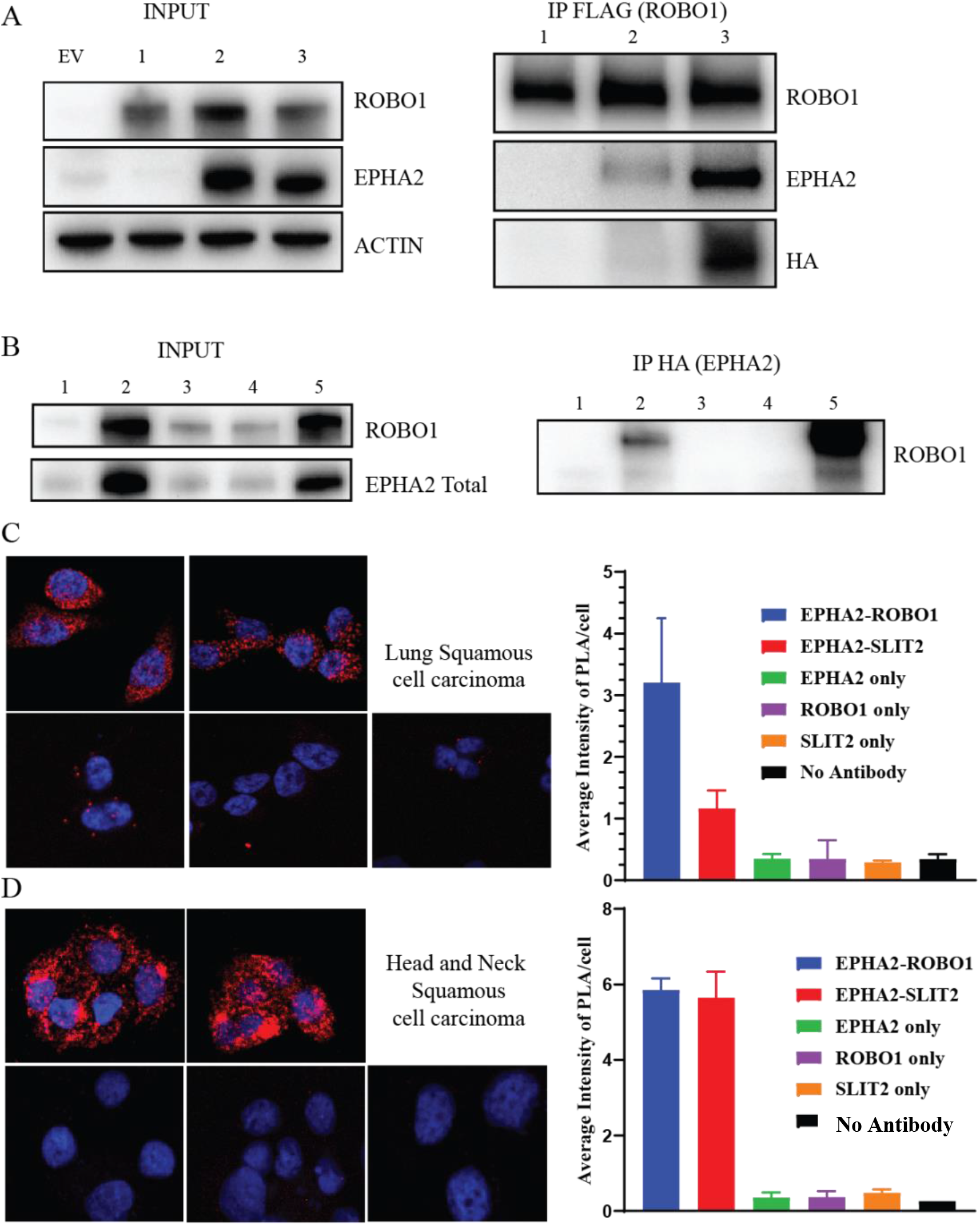
EPHA2-ROBO1 lies in the same protein complex shown by biochemical assays. Coimmunoprecipitation assay for EPHA2 and ROBO1. (A): Input and IP FLAG (ROBO1). EPHA2 wild type and EPHA2 kinase dead EPHA2(K645R KD) HA mutant is pulled down in complex with ROBO1. (EV: Empty vector (mCherry FLAG); 1: ROBO1-FLAG; 2: ROBO1-FLAG+EPHA2-WT-HA; 3: ROBO1-FLAG+EPHA2-(K645R KD)-HA) (B): Input and Inverse IP HA of EPHA2 pulls down WT ROBO1 but the phosphorylation mutants of ROBO1 do not bind to EPHA2. (EV: Empty vector (mCherry FLAG); 1: mCherry-FLAG+EPHA2-WT-HA; 2: ROBO1-FLAG+EPHA2-WT-HA; 3: ROBO1-FLAG Y932F+EPHA2-WT-HA; 4: ROBO1-FLAG Y1073F+EPHA2-WT-HA; 5: ROBO1-WT-FLAG+EPHA2-(K645R KD) HA) (C): Proximity Ligation Assay. Interaction of EPHA2-ROBO1 versus EPHA2-SLIT2 in Lung squamous cell carcinoma (SW900) and confocal microscopy quantification of punctate staining. (D): Proximity Ligation Assay. Interaction of EPHA2-ROBO1 versus EPHA2-SLIT2 in head and neck squamous cell carcinoma (SCC1) and confocal microscopy quantification of punctate staining.

Finally, we ascertained that there is an interaction between EPHA2-ROBO1 and EPHA2 and SLIT2 ligand of ROBO1, by performing a Proximity Ligation Assay (PLA) in both LSCC (SW900) and HNSCC (SCC1) cell lines. The results showed that EPHA2, ROBO1 and SLIT2 interact with each other in both the cell lines, however, the HNSCC showed more robust interactions between EPHA2/ROBO1 and EPHA2/SLIT2 compared to LSCC (**Fig. 5C & Fig. 5D**). These results suggest that EPHA2 and ROBO1 can form a heterodimer that may signal via SLIT2 to exert its effect on the tumor cells in both HNSCC and LSCC.

### SLIT2/ROBO1 interaction negatively regulates cellular proliferation and EPHA2 tumorigenic signaling

To assess the functional significance of the EPHA2/ROBO1 interaction, we examined the changes in EPHA2 signaling in the ROBO1 overexpressing NSCLC cells. Upon binding to the EphrinA1, EPHA2 induces autophosphorylation of Y772 at the activation loop and Y588, Y594 at its juxtamembrane region, leading to the recruitment of downstream effectors (*48, 15, 22*). Previous reports have also identified a ligand-independent activity of EPHA2 that is associated with phosphorylation at S897 by AKT and RSK (*23, 28*). Overexpression of ROBO1 in NSCLC cells resulted in down regulation of pS897, the ligand-independent activity marker of EPHA2. A more pronounced down regulation of pS897 signal was observed when cells (SK-MES-1 and H2170) were treated with SLIT2 (**Fig. 6A**). The western blot was quantified using densitometry plot (**Fig. 6B**). When cells were treated with EphrinA1, as expected, it enhanced ligand-dependent tumor suppressive signaling via pY588 and suppressed tumorigenic signaling via PS897 (**Fig. 6C**). Further, addition of EGF to these cells not only activated EGF receptor and its downstream effectors like AKT but also enhanced the tumorigenic signature of Phospho-EPHA2 S897 (**Fig. 6D**).

**Figure 6.**
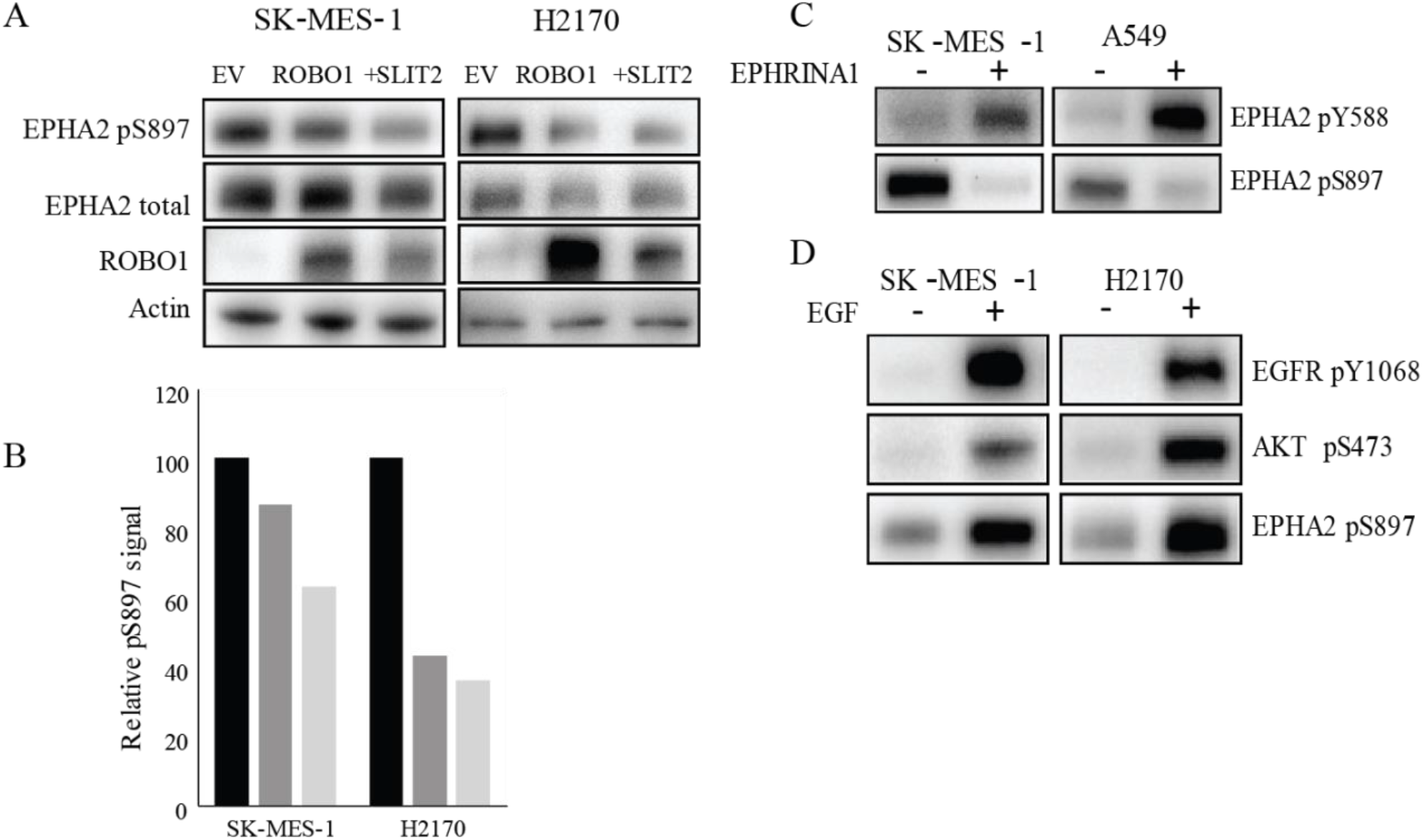
EPHA2-ROBO1 and EGF pathway cross talk. (A): Lung squamous cancer cell lines (SK-MES-1; H2170), ROBO1 was overexpressed and SLIT2(2ug/ml) treatment was done to look at total EPHA2 and ROBO1 levels and P-S897 levels. (B): Densitometry of the bands of western blot in (A) normalized to band intensity of empty vector. (C): EphrinA1 treatment was done in SK-MES-1 and A549 cells to look at canonical signaling (P-Y588) and non-canonical signaling (P-S897) signatures. (D): EGF treatment to SK-MES-1 and H2170 cells to assay P-S897 levels and activating phosphorylation of AKT and EGFR.

To assess the physiological changes in response to these ligands, we determined the effects of individual and combined treatments on cell proliferation and caspase activity. The cells were tagged with m-Kate2 (nuclear red fluorescence) and their proliferation and caspase activity were monitored in real time using the IncuCyte Live Cell Imaging System. In SW900 LSCC cell lines with high EPHA2 and low ROBO1 expression (majority patient cases based on TMA), the treatment with SLIT2 resulted in 34% inhibition of cellular proliferation. This inhibition was rescued by sequential treatment with EGF (**Fig. 7A**). Consistent with this finding, we observed that EGF leads to activation of PS897 of EPHA2 via AKT (**Fig. 6D**). Inhibition of cellular growth was also observed with the EPHA2 inhibitor ALW-II-41-27 (54%) and the combination treatment with SLIT2 showed an additive effect (61%) (**Fig. 7B**). Therefore, combination treatment will be most beneficial for LSCC with high EPHA2 and low ROBO1 levels. A similar inhibition of cell proliferation was observed in HNSCC (SCC1) with high EPHA2 and low ROBO1 expression, biggest fraction of cases with this type of cancer. Addition of SLIT2 resulted in 23% inhibition of cell proliferation that was rescued by the addition of EGF. But in SCC1 cell line ALW-II-41-27 alone and in combination with SLIT2 both showed proliferation inhibition of around 20%. Combination treatment did not have an additive effect (**Fig. 7A and B**). Thus, the HNSCC patients with high EPHA2 and low ROBO1, could benefit from treatment with either SLIT2 or Ensartinib. We also performed these experiments with another EPHA2 inhibitor Ensartinib (small molecule inhibitor for receptor tyrosine kinase that inhibits EPHA2), used in clinical settings. Ensartinib showed a similar effect on LSCC and HNSCC cell lines as observed with ALW-II-41-27 inhibitor. Ensartinib treatment in the LSCC SW900 resulted in 41% cellular inhibition. Combination treatment of Ensartinib and SLIT2 in LSCC(SW900) and HNSCC (SCC1) cell lines showed 65% and 70% inhibition respectively (**Fig. 7C and D**). Together, these results suggested that the EPHA2 and ROBO1/SLIT2 pathway crosstalk to inhibit cell proliferation.

**Figure 7.**
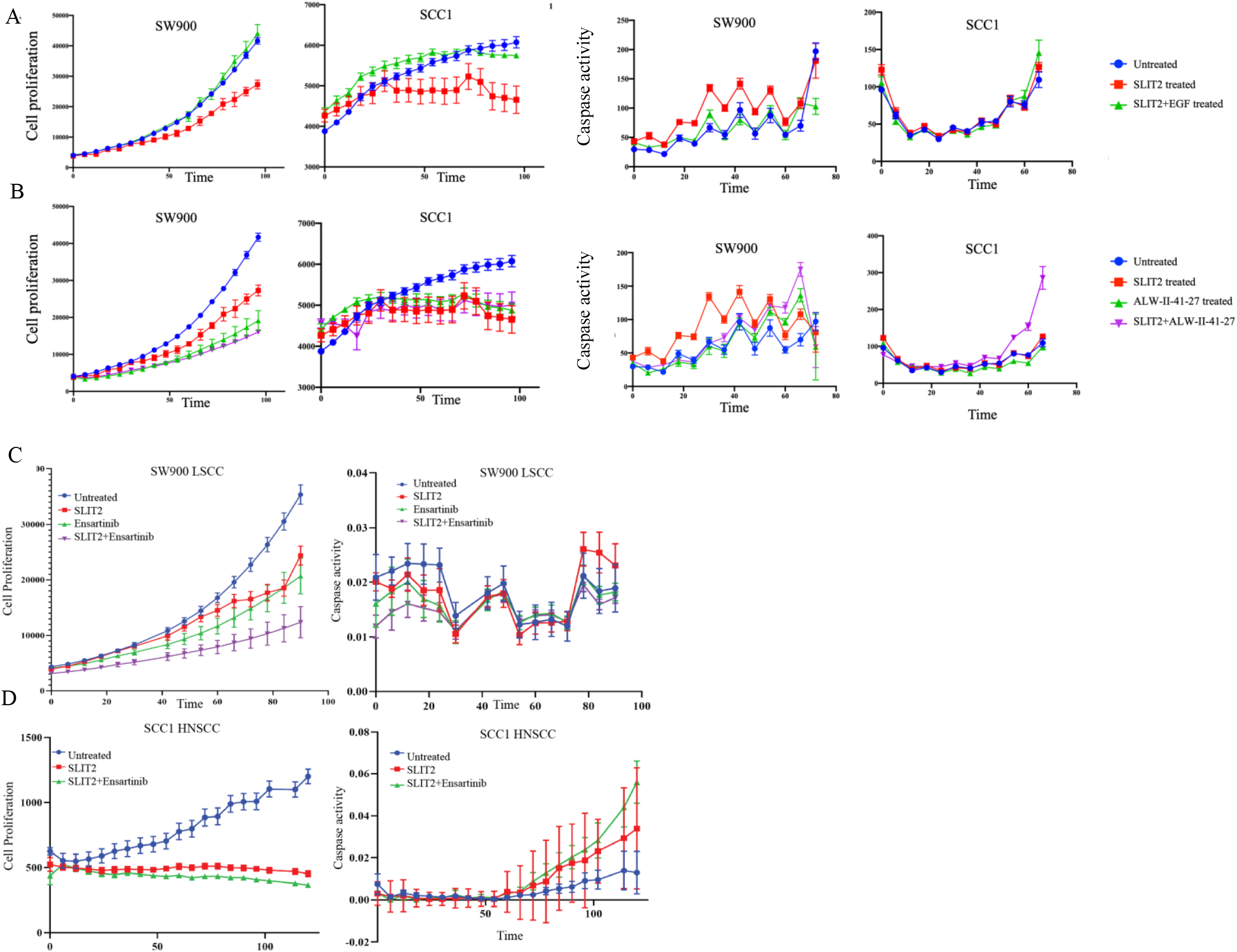
LSCC (SW900) and HNSCC (SCC1) cell line treatments of SLIT2, EGF, ALW-II-41-27 and Ensartinib to study cellular proliferation and caspase activity. (A) Incured SW900 lung squamous cancer cell line and SCC1, HNSCC cell line proliferation plot and caspase activity plot after treatment with SLIT2 (2ug/ml) and EGF. (B) Incured SW900 lung squamous cancer cell line and SCC1, HNSCC cell line proliferation plot and caspase activity plot after treatment with SLIT2 (2ug/ml) and ALW-II-41-27 (2uM).(C) Incured SW900 lung squamous cancer cell line proliferation plot and caspase activity plot after treatment with SLIT2 (2ug/ml), Ensartinib 3.461uM and SLIT2+Ensartinib combination treatment, (D) Incured: SCC1 head and neck squamous cancer cell line, proliferation plot and caspase activity plot after treatment with SLIT2 (2ug/ml) and SLIT2+Ensartinib 1.794uM combination treatment.

### Clinical significance of ROBO1 and EPHA2 expression

To investigate the potential roles of different ROBOs in NSCLC, particularly lung squamous cell carcinoma, we performed correlation studies using publicly available data from The Cancer Genome Atlas (TCGA) and cBioPortal. Of the four ROBO genes in humans, ROBO1 and ROBO2 showed decreased copy numbers in lung squamous cell carcinoma when compared to normal lung tissue (**Supplementary Figs. 3,4**). ROBO1 mutations showed a highly statistically significant association with survival (p = 8.373e-4), while the other ROBOs showed p value >0.05 (**Supplementary Fig. 2B**). In addition, more than half of the alterations of ROBO1 and ROBO2 genes in NSCLC samples were deletions and only a small percentage of cases of amplification were observed (**Supplementary Fig. 2C)**. Taken together, these data suggested that ROBO1 may act as a tumor suppressor and potentially antagonize EPHA2 in NSCLC.

### EPHA2, ROBO1 and SLIT form a complex

The STRING database (https://string-db.org/cgi/network.pl) is a protein interaction database that catalogs known and predicted protein-protein interactions. The interactions reported here include direct (physical) and indirect (functional) associations. These interactions stem from computational prediction, knowledge transfer between organisms, and interactions aggregated from other (primary) databases. A query of the STRING database revealed that EPHA2 and SLIT1 can interact directly and this interaction is experimentally validated in mouse and yeast (**Fig. 8A**). As far as we are aware, the interaction of EPHA2 and ROBO1, first shown in a comprehensive study done in *C. elegans,* has not been reported in humans as yet. In *C. elegans*, it was shown that there is a physical interaction between the VAB-1(EPH) tyrosine kinase domain binding to the intracellular portion of SAX-3(ROBO), specifically at the juxtamembrane and CC1 (conserved cytoplasmic region 1) regions. Therefore, based on the co-expression of EPHA2 and ROBO1/SLIT2 and the fact that they physically interact, we conclude that the EPHA2/ROBO1 heterodimer is functional and responds to the SLIT2 (**Fig. 8B**). This interaction blocks the phosphorylation of S897 as seen by a reduction in the levels of P-S897 which, in turn, attenuates cellular proliferation (tumorigenesis).

**Figure 8.**
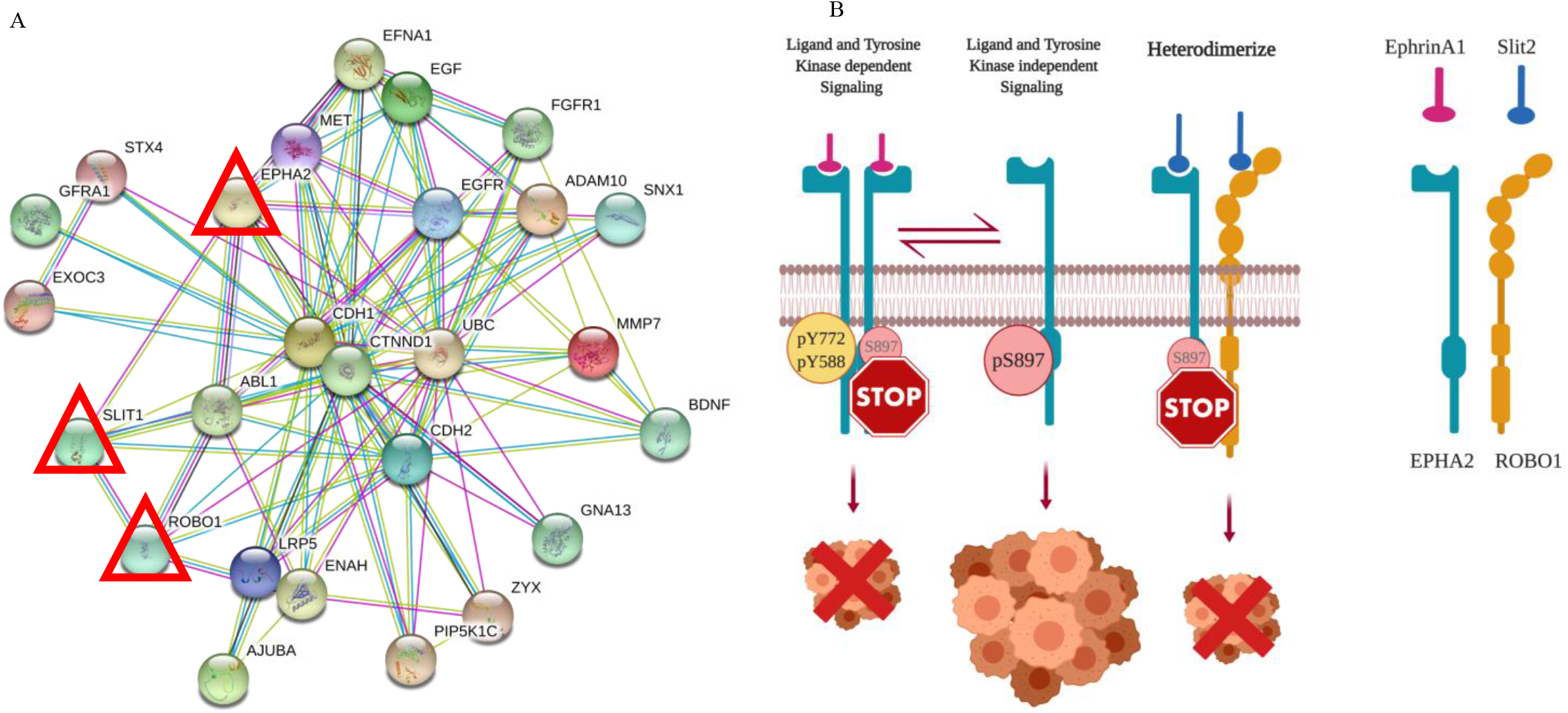
STRING database cross talk between EPHA2-ROBO1 and SLIT pathways and EPHA2-ROBO1 signaling model. (A): STRING database shows direct interaction between EPHA2 and SLIT1 with experimental evidence in other species like mouse, yeast. EPHA2-ROBO1 direct interaction has not been shown yet but the two proteins lie in the same network. (B). Model of EPHA2-ROBO1 heterodimerization being stabilized in presence of SLIT2 and in turn promoting non-tumorigenic signaling yielding therapeutic advantage.

## Discussion

Receptor tyrosine kinases that play critical roles in several biological processes have complex signaling mechanisms. Upon binding to their ligands, they often oligomerize and cross-phosphorylate each other, especially on tyrosine residues at the critical juxtamembrane domain and at the activation loop (*49–51*) to transduce various cellular signals. In the case of the EPH receptors, many of which are upregulated in numerous cancers, ligand activation leads to inhibition of tumor growth, migration and invasiveness (*51*). In our study when SCC cells were treated with EphrinA1 the natural ligand of EPHA2 (non-tumorigenic signaling) in *in vitro* culture, we did not see an inhibition in cellular proliferation of LSCC and HNSCC cells underscoring the need to study the presence of other degenerate unexplored pathways regulating EPHA2 signaling. Cancers overexpressing EPHA2 are accompanied by a concomitant loss of its ligands, implying that EPHA2 may act as an oncogene in absence of its ligand. Consistent with these observations, several studies have demonstrated that EPHA2 promotes cell migration and tumor malignancy in a ligand-independent manner. This signaling is characterized by high levels of S897 phosphorylation and low levels of Y772 and Y588 phosphorylation of EPHA2 (*23, 25–27*). In addition to ligand-independent activation, RTK heterodimerization or alternative ligands binding to atypical receptors is well documented and is recognized as a means to amplify signal or induce functional diversity (*52, 53*). For example, erythropoietin which is known to stimulate tumor growth by binding to EPHB4 that is not its natural receptor (*54*).

A physical interaction between the tyrosine kinase domain-1 VAB-1/EPHR and the CC1 region of SAX-3/ROBO has been observed in *C. elegans* (*31*). However, to the best of our knowledge, a functional heterodimer between EPHA2 and ROBO1 that can respond to SLIT2 signaling and attenuate cell proliferation in the human has not been reported thus far. We also believe that binding of SLIT2 to EPHA2, observed in the present study, has not been reported previously.

Studies by Dunaway et.al. 2011(*55*) have shown that cooperation between SLIT2 and EphrinA1 regulates a balance between the pro- and antiangiogenic functions of SLIT2 in mouse endothelial cells. However, no direct binding of SLIT2 to the EPHA2 receptor was shown. Nonetheless, querying the STRING protein interaction database showed evidence for physical interaction between EPHA2-SLIT1 in mouse and yeast lending further credence to our observations.

To summarize our findings, we propose a model (**Fig. 8B**) wherein, ROBO1 along with SLIT2 exerts its tumor-suppressive function by heterodimerizing with EPHA2 and blocking EGF induced phosphorylation of EPHA2 at S897 phosphorylation possibly by inhibiting AKT activity. In the LSCC cell line SW900 and HNSCC cell line SCC1 that expresses low levels of ROBO1, addition of SLIT2 inhibited cell proliferation (**Fig. 7A**), suggesting that in addition to binding to the EPHA2 and ROBO1 heterodimer (**Fig. 5C and D**) it may also bind to EPHA2 homodimer to exert its tumor suppressor effects. Alternatively, EPHA2 may heterodimerize with another ROBO receptor in low ROBO1-expressing cells to inhibit cell proliferation. EPHA2 over-expression in SCCs is significantly correlated with tumor site, T classification, clinical stage, recurrence, and lymph node metastasis (*56*). Therefore, we believe the results from the present study provide a new perspective on the treatments for LSCC and HNSCC patients who currently have limited options especially those who have acquired resistance to EGFR inhibitors or have progressed on platinum-based therapy. Thus, these patients can be stratified based on the expression patterns of EPHA2 and ROBO1 proteins in the diseased tissue (**Patient stratification Fig. 9**). The vast majority of LSCC patients(85%) with high EPHA2 and low ROBO1 expression can be treated with combination treatment of SLIT2 and Ensartinib due to additive effect of the two, while those patients with high EPHA2 and low ROBO1 expression in HNSCC (55%) like SCC1 can be treated with either SLIT2 or Ensartinib. Since EPHA2 and ROBO1 serve as biomarkers for discerning patients for appropriate therapy as well as therapeutic targets, they represent novel theranostics for these two diseases. The patient stratification presented here is based on EPHA2 and ROBO1 expression in TMAs. Performing the cell proliferation inhibition experiments with additional LSCC and HNSCC cell lines with varying degree of EPHA2 and ROBO1 expression will further validate these stratification strategies.

**Figure 9.**
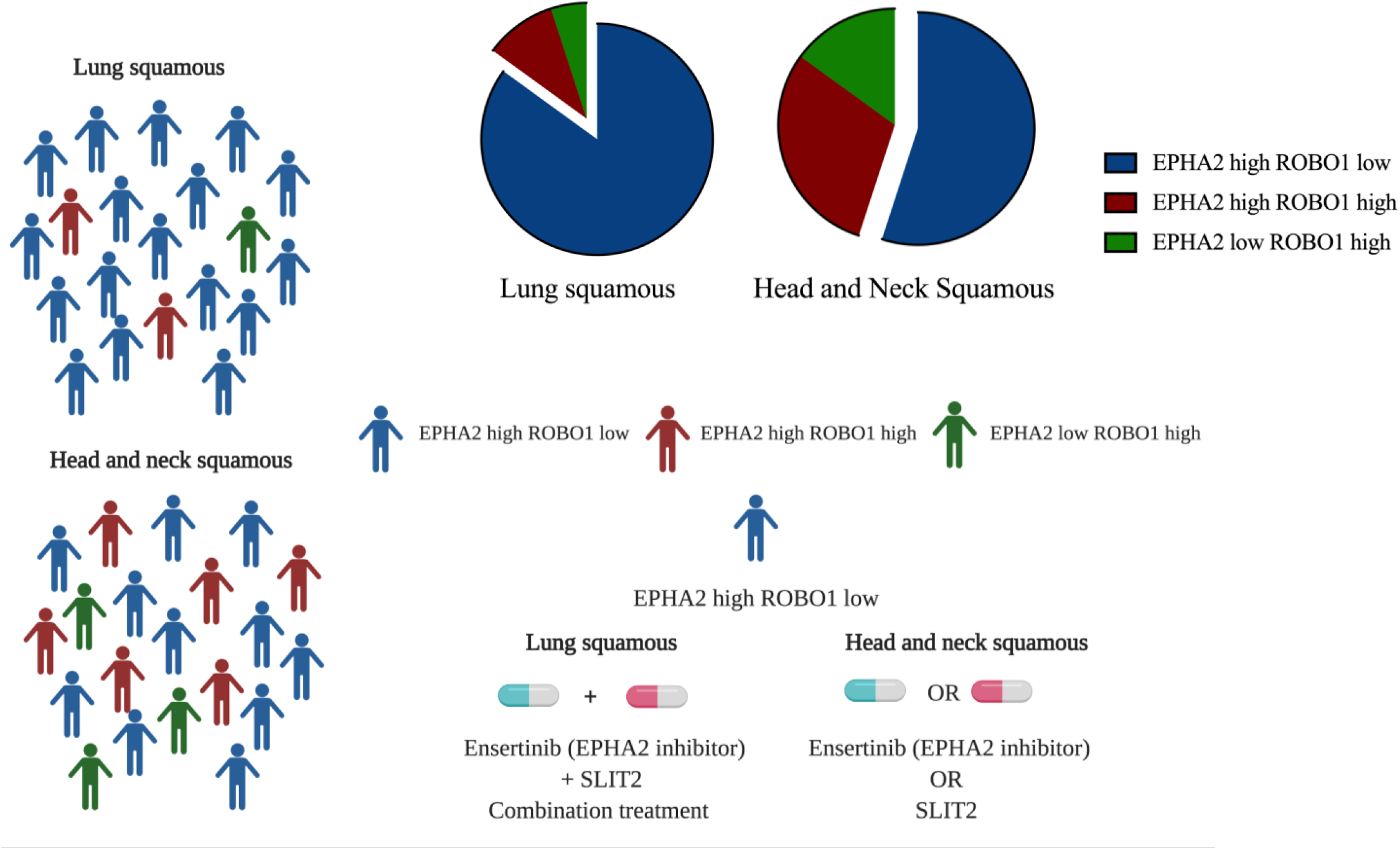
Patient stratification figure, treatment strategy for high EPHA2 and low ROBO1 patients in LSCC and HNSCC.

### Limitations of this study

Although the demonstration that EPHA2 and ROBO1 interact by multiple techniques is very appealing, and the cross-functional activation of this heterodimer by SLIT2 is novel, one of the limitations of the present study is that it does not provide information on the contacts made between the proteins that could aid small molecule therapeutic development or the development of novel SLIT2 mimetics that interact with the heterodimer more avidly. Additional studies in the future with more sophisticated biophysical techniques such as solution NMR or X-ray crystallography can provide deeper insight.

## Materials and Methods

### Cell Culture and Reagents

All NSCLC cell lines were from American Type Culture Collection (ATCC) (Manassas, VA, USA). NSCLC cell lines A549, H1993, SK-LU-1, H226, H1703, H2170, SK-MES-1, SW900 and the nonmalignant and immortalized control cell line BEAS-2B, were cultured in RPMI 1640 medium (Gibco/BRL) supplemented with 10% (v/v) fetal bovine serum (FBS), L-glutamine and 1% penicillin-streptomycin. HEK293 cells were cultured in DMEM medium (Gibco/BRL) supplemented with 10% (v/v) fetal bovine serum (FBS), L-glutamine and 1% penicillinstreptomycin. All HNSCC cell lines were obtained from indicated sources (supplementary table 1) All cell lines were cultured at 37°C with 5% CO_2_. ALW-II-41-27 was purchased from MedChemExpress (Monmouth Junction, NJ 08852, USA). EphrinA1, soluble EPHA2 and SLIT2 were purchased from R&D Systems (Minneapolis, MN 55413, USA). EGF ligand was purchased from Stem cell technology.

### C. elegans RNAi and drug treatment

Culture and handling of *C. elegans* were carried out as described (*57*). *vab-1* mutants (*OK1699, e2 and e2047*) and sax-3 mutants (*ky200 and ky123*) were obtained from *Caenorhabditis* Genetic Center. RNAi knock down was carried out by bacterial feeding method (*58*). Single wild type or mutant L3 worm were placed onto L4440 E. coli expressing either no RNA, or dsRNA targeting *vab-1* or *sax-3*. Total number of viable and dead F1 embryos were scored. For treatment of ALW-II-41-27, the indicated amount of drug or DMSO was added to 0.5 ml of base agar in 12-well plates and allowed to diffuse for 2 h. Feeding bacteria OP50 were then added on top of agar. A single L4 worm was then placed into an individual well. Viable F1 worms were then scored.

### siRNA, shRNA, DNA vectors and transfection

shRNA plasmids were constructed by inserting annealed oligonucleotide pairs targeting EPHA2, ROBO1 or luciferase into pJR288 as described (*59*). Nontargeting control pool siRNA (catalog no. D-001810) and SMARTpool siRNA targeting EPHA2 (catalog no. L-003116) were purchased from Dharmacon, Inc and utilized for transfection of HNSCC cell lines with a final concentration of 30 nmol/L siRNA with Lipofectamine RNAiMAX (Life Technologies). ROBO1 or EPHA2 expression vectors were constructed by fusing PCR fragments containing full length ROBO1 or EPHA2 upstream to eGFP, mCherry, mClover3, mRuby3, HA or FLAG sequences. All expression vectors were based on peGFP-N3 (Clontech/Takara, Mountain View, CA 94043, USA) with CMV promoter replaced by a eF1a promoter and eGFP replaced by mCherry, mClover3 or mRuby3. Point mutations were introduced by Q5 site directed mutagenesis kit according to manufacture protocol (NEB, Ispwich, MA 01938, USA). All constructs were confirmed by DNA sequencing. Oligonucleotides used are listed in supplementary table 2. All transfections were carried out using JetPrime transfection reagent according to manufacturer protocol (Polyplus transfection, 67400 Illkirch, France).

### Western blotting and Immunoprecipitation

Cell lysates for western blot were prepared by scraping cells and lysing them using RIPA buffer. Lysates were run on 4–15% or 4-20% Mini-protean TGX gels (Bio-Rad) and transferred onto ImmobilonTM membranes (MilliporeSigma, Burlingon, MA 01803, USA) or Turboblot system (Bio-Rad). Blots were blocked using 5% nonfat dry milk in TBST for 1 h and incubated with primary antibodies (listed in supplementary table 3) overnight at 4°C. After washing 3 times in TBST, blots were incubated with HRP-conjugated secondary antibodies for 1 h at room temperature. The blots were then washed three times and immuno-reactive bands were detected by WesternBright ECL (Advansta, San Jose, CA 95131, USA) or Azure Radiance (Azure) and imaged with ChemiDoc MP Imager (Bio-Rad) or Azure C600 (Azure). For coimmunoprecipitation assay, plasmids expressing EPHA2-HA and ROBO1-FLAG were cotransfected into HEK293 cells. Cells were collected 48 hours post-transfection and lysed by IP buffer (PBS + 1% triton with HALT protease and phosphatase inhibitor cocktails) (ThermoFisher, Waltham, MA 02451, USA). Lysates were adjusted to 1 mg/ml by IP buffer and protein complexes were immunoprecipitated by anti-FLAG magnetic beads (MilliporeSigma) or anti-HA magnetic beads (Thermofisher) at 40C for 4 h. Immunoprecipitated complexes were detected using a Western blot assay.

### Cell viability assays

NSCLC and HNSCC were labelled with m-Kate2 (red fluorescence) and stable cell lines were generated using puromycin selection. Incured (IncuCyte NucLight Rapid Red Reagent) cells were seeded in 96 well plates for 24 hours, followed by ligand treatment. SLIT2 (2ug/ml), EGF (0.5ug/ml) Ensartinib and ALW-II-41-27 IC50 doses. Cells were imaged every 6 hours for 72-h and their proliferation rates and caspase activity were plotted. Cell counting Kit 8 (Dojindo Molecular Technologies, catalog no. CK04) was used to determine relative numbers of viable cells 72 hours after post transfection with siRNA-targeting EPHA2 (siEPHA2) in HNSCC.

### Immunohistochemistry, Immunofluorescence Staining and live cells microscopy

Human lung cancer tissue micro array (LC642) was purchased from Biomax, Inc (Rockville, MD 20855, USA). EPHA2 was stained with anti-EPHA2 antibody (C-20, Santa Cruz Biotechnology, Dallas, TX 75220, USA), 1:200 for 30 min, and ROBO1 was stained with anti-ROBO1 antibody (PA5-29917, Invitrogen), 1:200 for 30 min. Each pair of stained TMAs was registered in Visiopharm before exporting a down sampled image. In FIJI, color deconvolution was used to extract the DAB staining (as grayscale) from each aligned TMA image, followed by false coloring the stains red or green. Quantification of stained area was performed using QuPath 0.1.2. Stain vectors were first estimated using the control core, followed by tissue detection to exclude most empty space, and finally positive pixel detection to calculate the percentage of tissue area within a given core that stained above the DAB threshold for the protein of interest (*60*). Then the percentage positive area was plotted using prism software.

For immunofluorescence staining, cells were seeded into Lab-Tek II Chamber Slide (Thermofisher) or Number 1 cover slips in a 24-well plate for 24 to 48 h, then fixed by 1% formaldehyde in PBS for 20 min at room temperature, permeabilized by PBS containing 0.1% Tween and 0.25 % Triton X-100. After three washes with PBS, fixed cells were blocked with 5% FBS in PBS for 1 h at room temperature. Primary antibodies were then added and incubated overnight at 40C. Primary antibodies were removed, and the slides were washed 5 times with PBS. Alexa Flour 488 or Alexa Flour 547 conjugated secondary antibodies and Hoechst 33342 dye (ThermoFisher) for staining nuclei were then added and allowed to incubate for 2 hours at room temperature. The slides/coverslips were then washed five times with PBS and mounted in Prolong Gold Antifade reagent (ThermoFisher).

For live cell imaging, transfected cells were plated onto 35 mm Delta TPG dish (Bioptechs, Butler, PA16002, USA) for 24 h. The dishes were then placed on temperature controlled microscopic stage that was connected to CO_2_ supply. All images were acquired by Zeiss LSM880 confocal microscope and analyzed by Zen software (Zeiss USA, Thornwood, NY 10594, USA).

### Proximity Ligation Assay (PLA)

To perform a complete Duolink^®^ PLA in situ experiment we used two primary antibodies (PLA, Immunofluorescence validated) that recognize EPHA2, ROBO1 or SLIT2 epitopes. The starter kit from SIGMA supplies all other necessary reagents for Duolink^®^ PLA reactions, which include a pair of PLA probes (Anti-Rabbit PLUS and Anti-Mouse MINUS), red detection reagents, wash buffers, and mounting medium. The primary antibodies used came from the same species as the Duolink^®^ PLA probes for EPHA2/ROBO1 or EPHA2/SLIT2 PLA. Analysis was carried out using standard immunofluorescence assay technique. We used confocal microscope to capture images. For the quantification of this staining the confocal images were extracted, and the binary image was generated. The binary image was thresholded using FIJI software. The average intensity was measured and plotted to compare the binding of the two proteins assayed.

## Supporting information

Ka M. Pang et.al. Supplementary Movie

Ka M. Pang et. al. Supplementary materials

## Author contributions

Ka M. Pang: conceptualized, performed experiments, analyzed data, wrote and reviewed the paper; Saumya Srivastava: conceptualized, performed experiments, analyzed data, wrote and reviewed the paper; Mari Iida: performed experiments, analyzed data, wrote and reviewed the paper; Michael Nelson: analyzed data, wrote and reviewed the paper; Arin Nam: performed experiments, reviewed the paper; Jiayi Liu: analyzed data; Jiale Wang: performed experiments; Isa Mambetsariev: provided patient samples; Atish Mohanty: conceptualized and reviewed the paper; Nellie McDaniel: performed experiments, analyzed data; Amita Behal: conceptualized, wrote and reviewed the paper; Prakash Kulkarni: reviewed and wrote the paper; Deric L. Wheeler: conceptualized the idea, reviewed the paper; Ravi Salgia: conceptualized the idea, reviewed the paper

## Data Availability

All data are contained within the manuscript and supplementary video is attached separately.

## References

1. https://www.who.int/news-room/fact-sheets/detail/cancer.

2. https://www.wcrf.org/dietandcancer/lung-cancer.

3. Bray F, Ferlay J, Soerjomataram I, et al. Global cancer statistics 2018: GLOBOCAN estimates of incidence and mortality worldwide for 36 cancers in 185 countries. CA Cancer J Clin 2018; 68:394.

4. Siegel RL, Miller KD, Jemal A. Cancer statistics, 2020. CA Cancer J Clin 2020; 70:7.

5. Zappa C, Mousa SA. Non-small cell lung cancer: current treatment and future advances Transl Lung Cancer Res. 2016 Jun; 5(3):288–300.

6. Joshua K. Sabari and Paul K. Relevance of genetic alterations in squamous and small cell lung cancer. Ann Transl Med. 2017 Sep; 5(18): 373.

7. Thayer K. Darling and Tracey J. Lamb. Emerging Roles for Eph Receptors and Ephrin Ligands in Immunity. Front Immunol. 2019; 10: 1473.

8. Birgit Mosch, Bettina Reissenweber, Christin Neuber, and Jens Pietzsch. Eph Receptors and Ephrin Ligands: Important Players in Angiogenesis and Tumor Angiogenesis. J Oncol. 2010; 2010: 135285.

9. Arvinder Singh, Emily Winterbottom, and Ira O. Daar. Eph/ephrin signaling in cell-cell and cell-substrate adhesion. Front Biosci (Landmark Ed). 2012 Jan 1; 17: 473–497.

10. Erika M. Lisabeth, Giulia Falivelli, and Elena B. Pasquale. Eph Receptor Signaling and Ephrins Cold Spring Harb Perspect Biol. 2013 Sep; 5(9).

11. Zelinski DP1, Zantek ND, Stewart JC, Irizarry AR, Kinch MS. EPHA2 overexpression causes tumorigenesis of mammary epithelial cells. Cancer Res. 61, 2301–6 (2001).

12. Fang WB1, Brantley-Sieders DM, Parker MA, Reith AD, Chen J. A kinase-dependent role for EPHA2 receptor in promoting tumor growth and metastasis. Oncogene. 24, 7859–68 (2005).

13. Faoro L1, Singleton PA, Cervantes GM, Lennon FE, Choong NW, Kanteti R, Ferguson BD, Husain AN, Tretiakova MS, Ramnath N, Vokes EE, Salgia R. EPHA2 mutation in lung squamous cell carcinoma promotes increased cell survival, cell invasion, focal adhesions, and mammalian target of rapamycin activation. J Biol Chem. 285, 18575–85 (2010).

14. Ferguson BD1, Liu R, Rolle CE, Tan YH, Krasnoperov V, Kanteti R, Tretiakova MS, Cervantes GM, Hasina R, Hseu RD, Iafrate AJ, Karrison T, Ferguson MK, Husain AN, Faoro L, Vokes EE, Gill PS, Salgia R. The EPHB4 receptor tyrosine kinase promotes lung cancer growth: a potential novel therapeutic target. PLoS One. 8, 67668 (2013).

15. Elena B. Pasquale. EPH receptors and ephrins in cancer: bidirectional signaling and beyond. Nat Rev Cancer. 10, 165–180 (2010).

16. Salgia R, Kulkarni P. The Genetic/Non-genetic Duality of Drug ‘Resistance’ in Cancer. Trends Cancer. 4, 110–118 (2018).

17. Tan YC, Srivastava S, Won BM, Kanteti R, Arif Q, Husain AN, Li H, Vigneswaran WT, Pang KM, Kulkarni P, Sattler M, Vaidehi N, Mambetsariev I, Kindler HL, Wheeler DL, Salgia R. EPHA2 mutations with oncogenic characteristics in squamous cell lung cancer and malignant pleural mesothelioma. Oncogenesis. 8, 49 (2019).

18. Katherine R. Amato, Shan Wang, Andrew K. Hastings, Victoria M. Youngblood, Pranav R. Santapuram, Haiying Chen, Justin M. Cates, Daniel C. Colvin, Fei Ye, Dana M. Brantley-Sieders, Rebecca S. Cook, Li Tan, Nathanael S. Gray, and Jin Chen. Genetic and pharmacologic inhibition of EPHA2 promotes apoptosis in NSCLC. J Clin Invest. 2014 May 1; 124(5): 2037–2049.

19. Zhuang G, Brantley-Sieders DM, Vaught D, Yu J, Xie L, Wells S, Jackson D, Muraoka-Cook R, Arteaga C, Chen J. Elevation of receptor tyrosine kinase EPHA2 mediates resistance to trastuzumab therapy. Cancer Res. 70, 299–308 (2010).

20. Koch H, Busto ME, Kramer K, Médard G, Kuster B. Chemical Proteomics Uncovers EPHA2 as a Mechanism of Acquired Resistance to Small Molecule EGFR Kinase Inhibition. J Proteome Res. 14, 2617–25 (2015).

21. Amato KR, Wang S, Tan L, Hastings AK, Song W, Lovly CM, Meador CB, Ye F, Lu P, Balko JM, Colvin DC, Cates JM, Pao W, Gray NS, Chen J. EPHA2 Blockade Overcomes Acquired Resistance to EGFR Kinase Inhibitors in Lung Cancer. Cancer Res. 76, 305–18 (2016).

22. Boyd AW, Bartlett PF, Lackmann M. Therapeutic targeting of EPH receptors and their ligands. Nat Rev Drug Discov. 13, 39–62 (2014).

23. Miao H, Li DQ, Mukherjee A, Guo H, Petty A, Cutter J, Basilion JP, Sedor J, Wu J, Danielpour D, Sloan AE, Cohen ML, Wang B. EPHA2 mediates ligand-dependent inhibition and ligand-independent promotion of cell migration and invasion via a reciprocal regulatory loop with Akt. Cancer Cell. 16, 9–20 (2009).

24. Taddei ML, Parri M, Angelucci A, Onnis B, Bianchini F, Giannoni E, Raugei G, Calorini L, Rucci N, Teti A, Bologna M, Chiarugi P. Kinase-dependent and - independent roles of EPHA2 in the regulation of prostate cancer invasion and metastasis. Am J Pathol. 174,1492–503 (2009).

25. Beauchamp A, Debinski W. EPHs and ephrins in cancer: Ephrin-A1 signalling. Semin Cell Dev Biol. 23,109–15 (2012).

26. Brantley-Sieders DM. Clinical relevance of EPHs and ephrins in cancer: lessons from breast, colorectal, and lung cancer profiling. Semin Cell Dev Biol. 23, 102–8 (2012).

27. Zhou Y, Sakurai H. Emerging and Diverse Functions of the EPHA2 Noncanonical Pathway in Cancer Progression. Biol Pharm Bull. 40,1616–1624 (2017).

28. Zhou Y, Yamada N, Tanaka T, Hori T, Yokoyama S, Hayakawa Y, Yano S, Fukuoka J, Koizumi K, Saiki I, Sakurai H. Crucial roles of RSK in cell motility by catalysing serine phosphorylation of EPHA2. Nat Commun.; 6, 7679 (2015).

29. George, S.E., Simokat, K., Hardin, J., and Chisholm, A.D. (1998). The VAB-1 Eph receptor tyrosine kinase functions in neural and epithelial morphogenesis in C. elegans. Cell 92, 633–643.

30. Ian D Chin-Sang, Sean E George, Mei Ding, Sarah L Moseley, Andrew S Lynch, Andrew D Chisholm. The Ephrin VAB-2/EFN-1 Functions in Neuronal Signaling to Regulate Epidermal Morphogenesis in C. elegans, Articleļ Volume 99, Issue 7, P781–790, December 23, 1999.

31. Ghenea S, Boudreau JR, Lague NP, Chin-Sang ID. The VAB-1 EPH receptor tyrosine kinase and SAX-3/ROBO neuronal receptors function together during C. elegans embryonic morphogenesis. Development. 132, 3679–90 (2005).

32. Bernadskaya YY, Wallace A, Nguyen J, Mohler WA, Soto MC. UNC-40/DCC, SAX-3/Robo, and VAB-1/Eph polarize F-actin during embryonic morphogenesis by regulating the WAVE/SCAR actin nucleation complex. PLoS Genet. 2012;8.

33. Rothberg JM, Hartley DA, Walther Z, Artavanis-Tsakonas S. SLIT: an EGF-homologous locus of D. melanogaster involved in the development of the embryonic central nervous system. Cell. 23,1047–59 (1988).

34. Kidd T1, Bland KS, Goodman CS. SLIT is the midline repellent for the ROBO receptor in Drosophila. Cell. 19, 785–94 (1999).

35. Xian J, Clark KJ, Fordham R, Pannell R, Rabbitts TH, Rabbitts PH. Inadequate lung development and bronchial hyperplasia in mice with a targeted deletion in the Dutt1/ROBO1 gene. Proc Natl Acad Sci U S A. 18, 15062–6 (2001).

36. Grieshammer U, Le Ma, Plump AS, Wang F, Tessier-Lavigne M, Martin GR. SLIT2-mediated ROBO2 signaling restricts kidney induction to a single site. Dev Cell. 6, 709–17 (2004).

37. Bedell VM, Yeo SY, Park KW, Chung J, Seth P, Shivalingappa V, Zhao J, Obara T, Sukhatme VP, Drummond IA, Li DY, Ramchandran R. roundabout4 is essential for angiogenesis in vivo. Proc Natl Acad Sci U S A. 3, 6373–8 (2005).

38. Strickland P, Shin GC, Plump A, Tessier-Lavigne M, Hinck L. SLIT2 and netrin 1 act synergistically as adhesive cues to generate tubular bi-layers during ductal morphogenesis. Development. 133, 823–32 (2006).

39. Chen H, Zhang M, Tang S, London NR, Li DY, Zhang K. SLIT-ROBO signaling in ocular angiogenesis. Adv Exp Med Biol. 664, 457–63 (2010).

40. Blockus H, Chédotal A. SLIT-ROBO signaling. Development. 143, 3037–44 (2016).

41. Ballard MS1, Hinck L. A roundabout way to cancer. Adv Cancer Res. 114,187–235 (2012).

42. Huang Z, Wen P, Kong R, Cheng H, Zhang B, Quan C, Bian Z, Chen M, Zhang Z, Chen X, Du X, Liu J, Zhu L, Fushimi K, Hua D, Wu JY. USP33 mediates SLIT-ROBO signaling in inhibiting colorectal cancer cell migration. Int J Cancer. 136, 1792–802 (2015).

43. Gara RK, Kumari S, Ganju A, Yallapu MM, Jaggi M, Chauhan SC. SLIT/ROBO pathway: a promising therapeutic target for cancer. Drug Discov Today. 20, 156–64 (2015).

44. Maiti GP, Ghosh A, Mondal P, Ghosh S, Chakraborty J, Roy A, Roy Chowdhury S, Panda CK. Frequent inactivation of SLIT2 and ROBO1 signaling in head and neck lesions: clinical and prognostic implications. Oral Surg Oral Med Oral Pathol Oral Radiol. 119, 202–12 (2015).

45. Hohenester E. Structural insight into SLIT-ROBO signalling. Biochem Soc Trans. 36, 251–6 (2008).

46. Choi Y1, Syeda F, Walker JR, Finerty PJ Jr, Cuerrier D, Wojciechowski A, Liu Q, Dhe-Paganon S, Gray NS. Discovery and structural analysis of Eph receptor tyrosine kinase inhibitors. Bioorg Med Chem Lett. 2009 Aug 1;19(15):4467–70.

47. Benchun Miao, Zhenyu Ji, Li Tan, Michael Taylor, Jianming Zhang, Hwan Geun Choi, Dennie T. Frederick, Raj Kumar, Jennifer A. Wargo, Keith T. Flaherty, Nathanael S. Gray, and Hensin Tsao. EphA2 is a Mediator of Vemurafenib Resistance and a Novel Therapeutic Target in Melanoma. Cancer Discov. 2015 Mar; 5(3): 274–287.

48. WB FangDM Brantley-SiedersY HwangAJ HamJ Chen. Identification and functional analysis of phosphorylated tyrosine residues within EphA2 receptor tyrosine kinase (2008) Journal of Biological Chemistry 283:16017–16026.

49. Singh DR, Ahmed F, King C, Gupta N, Salotto M, Pasquale EB, Hristova K. EPHA2 Receptor Unliganded Dimers Suppress EPHA2 Pro-Tumorigenic Signaling. J Biol Chem. 290, 27271–9 (2015).

50. Deo R. Singh, Elena B. Pasquale, and Kalina Hristova, A Small Peptide Promotes EphA2 Kinase-Dependent Signaling by Stabilizing EphA2 Dimers. Biochim Biophys Acta. 2016 Sep; 1860(9): 1922–1928.

51. Singh DR, Kanvinde P, King C, Pasquale EB, Hristova K. The EPHA2 receptor is activated through induction of distinct, ligand-dependent oligomeric structures. Commun Biol. 22, 1:15 (2018).

52. Ichiro N. Maruyama. Mechanisms of Activation of Receptor Tyrosine Kinases: Monomers or Dimers. Cells. 2014 Jun; 3(2): 304–330.

53. Lemmon MA1, Schlessinger J. Cell signaling by receptor tyrosine kinases. Cell. 2010 Jun 25;141(7): 1117–34.

54. Pradeep S, Huang J, Mora EM, Nick AM, Cho MS, Wu SY, Noh K, Pecot CV, Rupaimoole R, Stein MA, Brock S, Wen Y, Xiong C, Gharpure K, Hansen JM, Nagaraja AS, Previs RA, Vivas-Mejia P, Han HD, Hu W, Mangala LS, Z and B, Stagg LJ, Ladbury JE, Ozpolat B, Alpay SN, Nishimura M, Stone RL, Matsuo K, Armaiz-Peña GN, Dalton HJ, Danes C, Goodman B, Rodriguez-Aguayo C, Kruger C, Schneider A, Haghpeykar S, Jaladurgam P, Hung MC, Coleman RL, Liu J, Li C, Urbauer D, Lopez-Berestein G, Jackson DB, Sood AK. Erythropoietin Stimulates Tumor Growth via EPHB4. Cancer Cell. 28, 610–622. (2015).

55. Dunaway CM, Hwang Y, Lindsley CW, Cook RS, Wu JY, Boothby M, Chen J, Brantley-Sieders DM. Cooperative signaling between Slit2 and Ephrin-A1 regulates a balance between angiogenesis and angiostasis. Mol Cell Biol. 2011 Feb;31(3):404–16.

56. Liu Y1, Yu C, Qiu Y, Huang D, Zhou X, Zhang X, Tian Y. Downregulation of EphA2 expression suppresses the growth and metastasis in squamous-cell carcinoma of the head and neck in vitro and in vivo. J Cancer Res Clin Oncol. 2012 Feb;138(2):195–202.

57. Theresa Stiernagle. Caenorhabditis Genetics Center, University of Minnesota, Minneapolis, MN 55455 USA. Maintenance of C. elegans. Worm book the online review of C. elegans biology.

58. Kamath RS, Martinez-Campos M, Zipperlen P, Fraser AG, Ahringer J. Effectiveness of specific RNA-mediated interference through ingested double-stranded RNA in Caenorhabditis elegans. Genome Biol. 2001;2(1): RESEARCH0002. Epub 2000 Dec 20.

59. Ka Ming Pang, Daniela Castanotto, Haitang Li, Lisa Scherer, and John J Rossi. Incorporation of aptamers in the terminal loop of shRNAs yields an effective and novel combinatorial targeting strategy. Nucleic Acids Res. 2018 Jan 9; 46(1).

60. Peter Bankhead, Maurice B. Loughrey, José A. Fernández, Yvonne Dombrowski, Darragh G. McArt, Philip D. Dunne, Stephen McQuaid, Ronan T. Gray, Liam J. Murray, Helen G. Coleman, Jacqueline A. James, Manuel Salto-Tellez & Peter W. Hamilton QuPath: Open source software for digital pathology image analysis. Scientific Reports volume 7, Article number: 16878 (2017).

61. Kimple RJ, Smith MA, Blitzer GC, Torres AD, Martin JA, Yang RZ, et al. Enhanced radiation sensitivity in HPV-positive head and neck cancer. Cancer Res. 2013;73:4791–800.

62. Brenner JC, Graham MP, Kumar B, Saunders LM, Kupfer R, Lyons RH, et al. Genotyping of 73 UM-SCC head and neck squamous cell carcinoma cell lines. Head Neck. 2010;32:417–26.

