## Supplementary material for "The Ephrin receptor A2 and Roundabout Guidance Receptor 1 heterodimer: A potential theranostic for squamous cell carcinomas": Ka M. Pang et. al. Supplementary materials

**Supplementary figure 1:** *C. elegans* model: synthetic lethality of VAB-1 and SAX-3 pathways: (A & B): RNA interference experiment of VAB-1(EPHA2) and SAX-3 (ROBO1) to look at embryonic lethality in comparison to wild type worms N2. (C): The synthetic lethal phenotype is also seen with SAX-3 RNAi and use of a EPHA2 inhibitor ALW-II-41-27.

**Supplementary figure 2:** (A): Survival plot for lung cancer cases with high ROBO1 versus low ROBO1 expression. The statistical data is shown in Table 1. Patients with high ROBO1 expression have better survival. (B): Survival plot of cancer cases with alteration in ROBO1 gene versus cases with no ROBO1 gene alterations. The number of cases are represented in Table 2. (C): TCGA data of ROBO1 alteration frequency in lung squamous cancers.The plot represents more alterations are present in ROBO1 gene than ROBO2 gene in NSCLC samples.

**Supplementary figure 3:**TCGA data ROBO family members copy number (A-ROBO1, B-ROBO2, copy number in lung squamous cell carcinoma).

**Supplementary figure 4:** TCGA data ROBO family members copy number (A-ROBO3, B-ROBO4, copy number in lung squamous cell carcinoma).

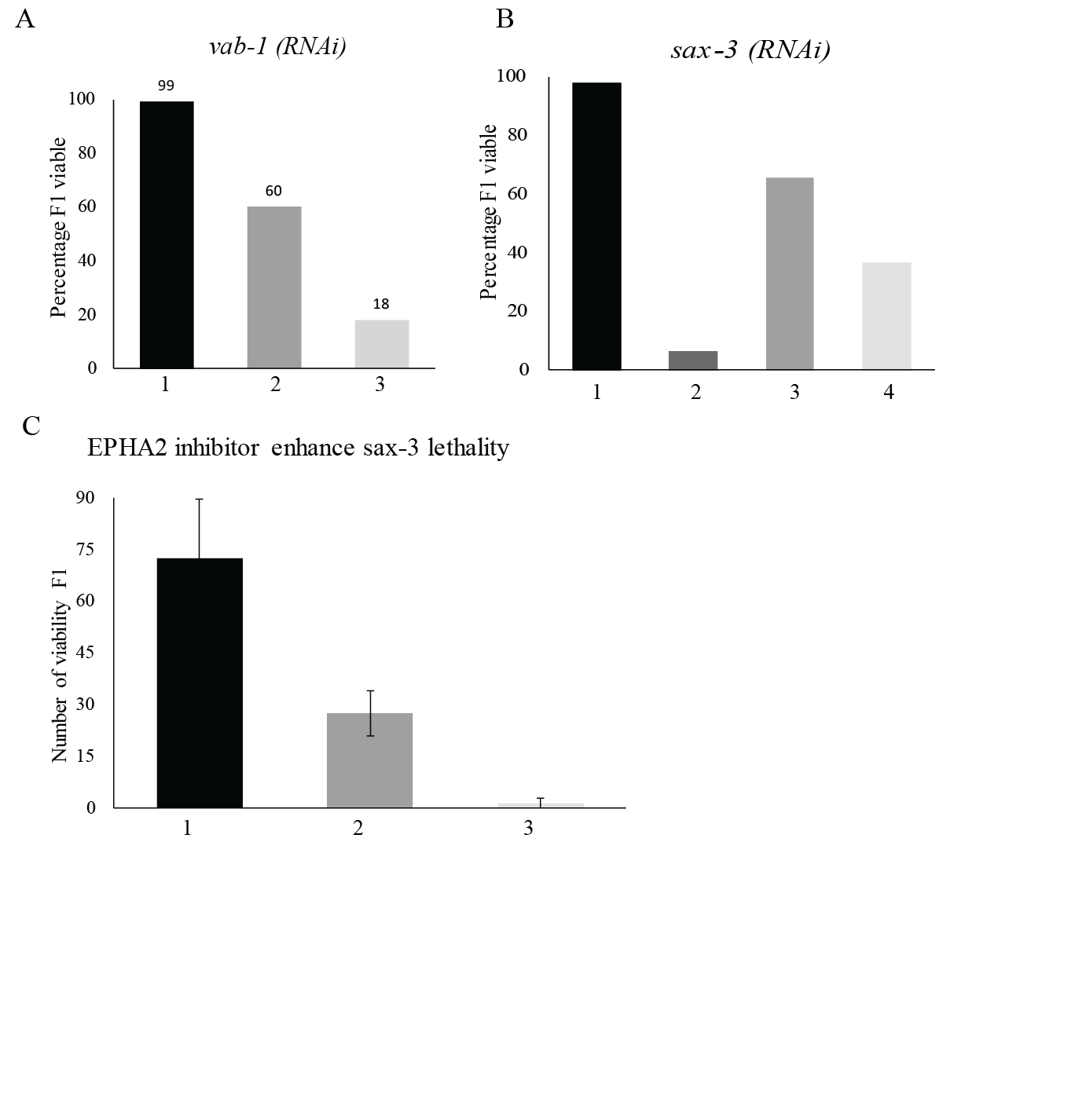

Number of viable F1

Supplementary figure 1

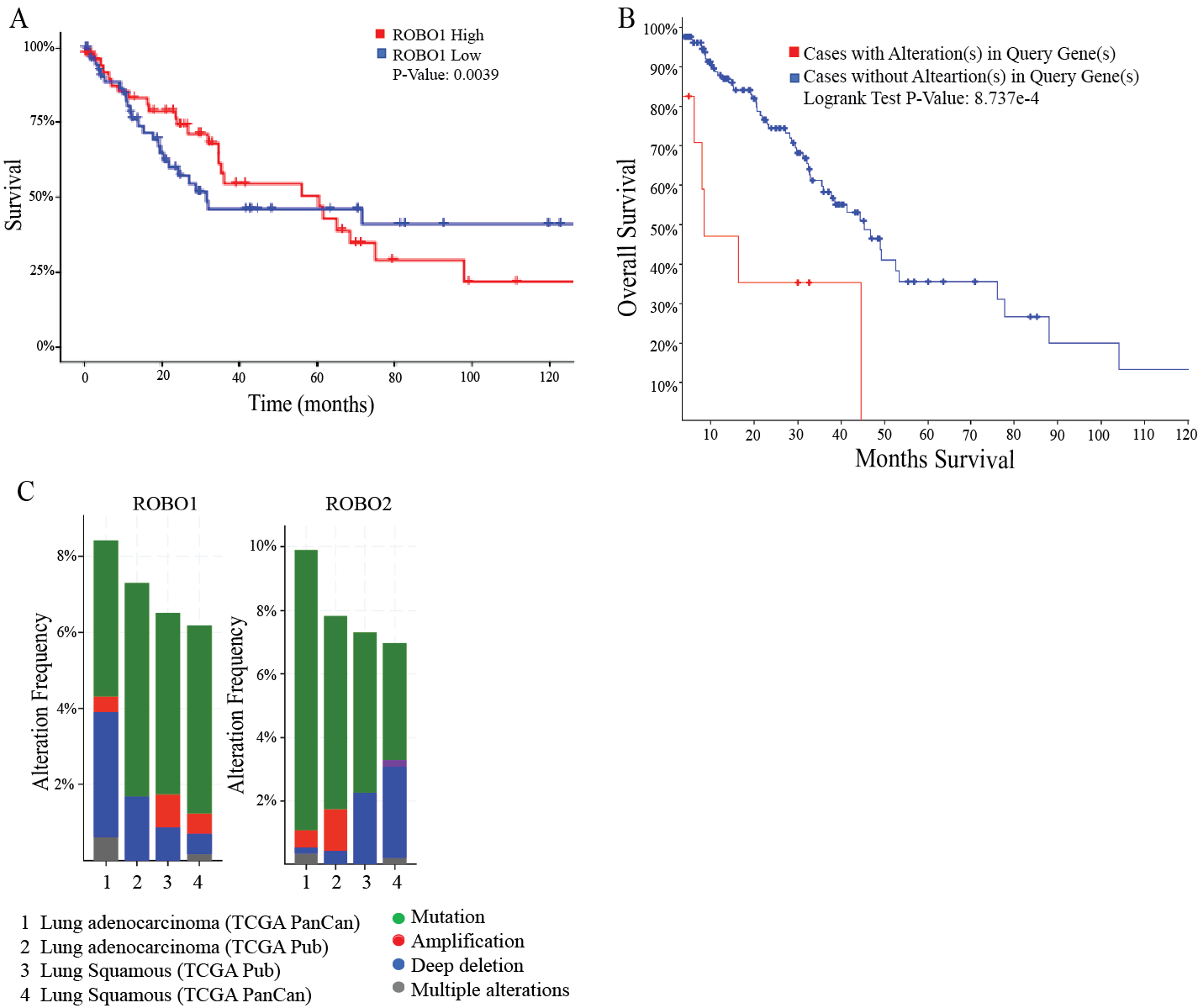

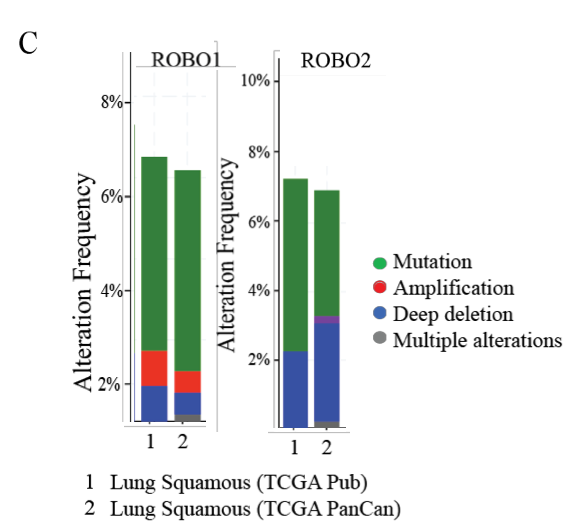

Supplementary figure 2

Supplementary figure 3

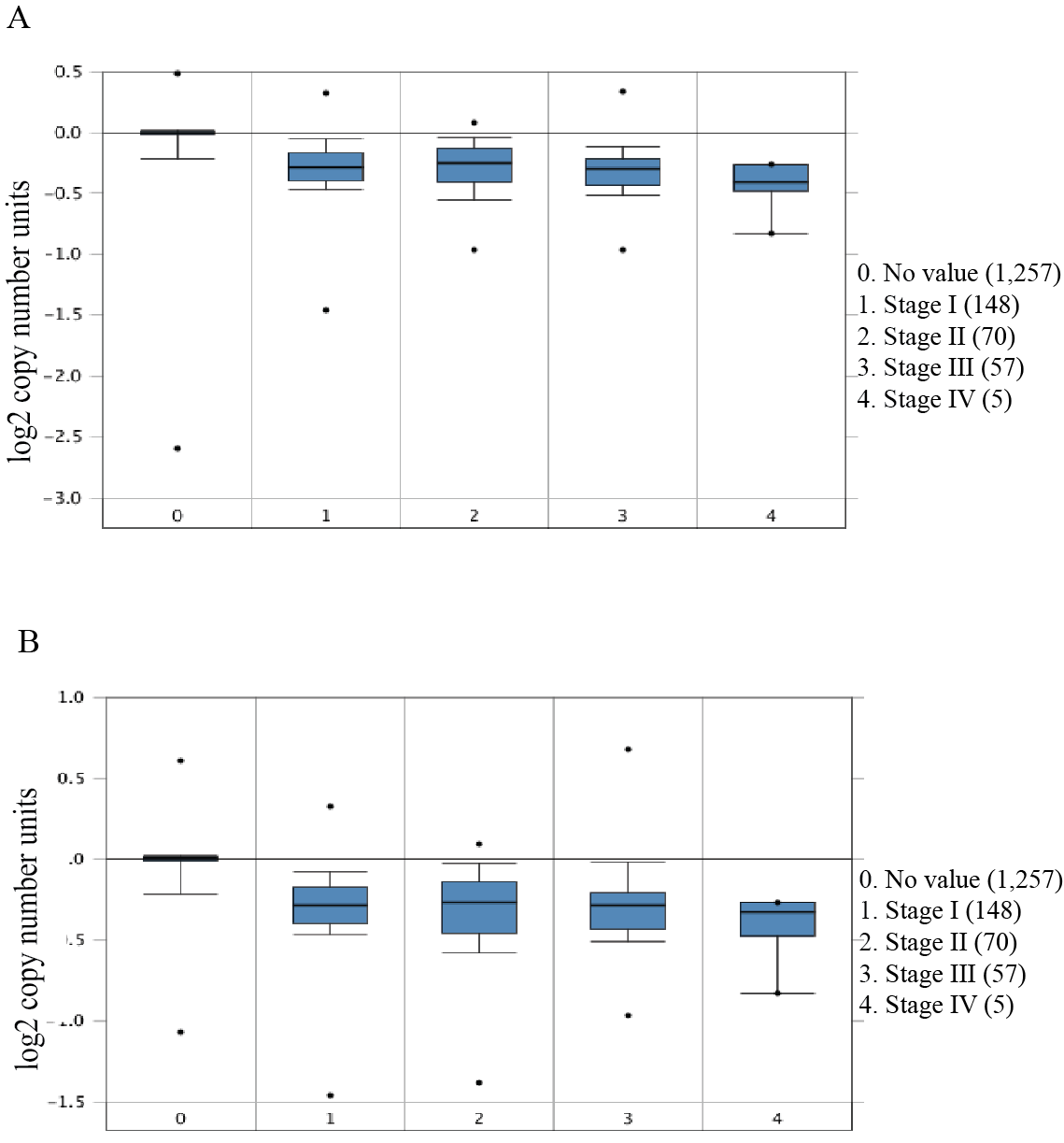

Supplementary figure 4

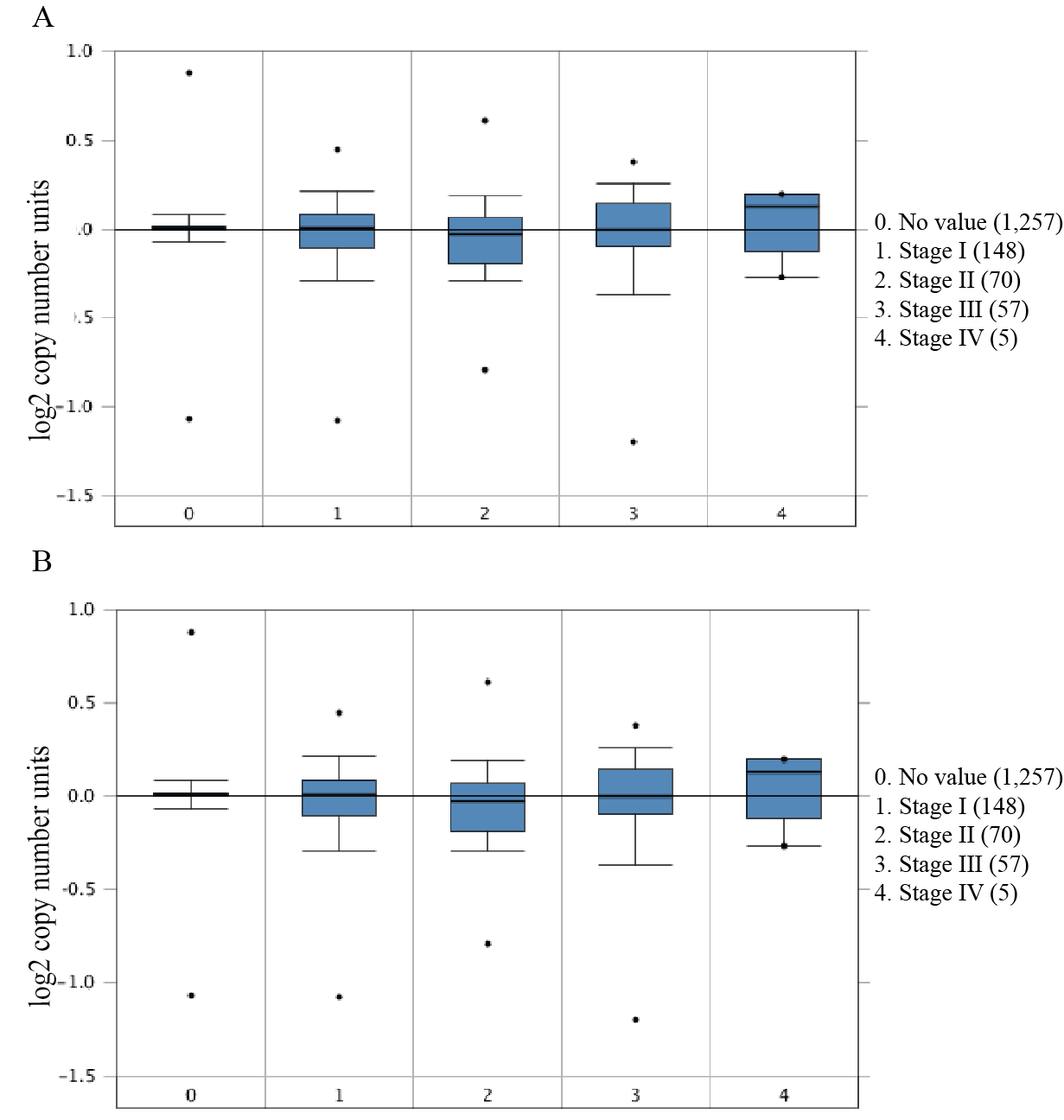

Supplementary tables

Supplementary table 1: Cell Lines*.* All HNSCC cell lines were obtained and validated by the following sources. HPV status of the following cell lines has been previously validated (*59, 60*).

| **Cell line** | **Sources** | **Culture condition** |
| --- | --- | --- |
| SCC-15  SCC-25 | American Type Culture Collection | DMEM/F12 (1:1), 10% FBS, 400ng/ml, hydrocortisone, penicillin (100 units/mL), streptomycin (100 mg/mL) |
| 93-vu-147T | Dr. Robert Ferris, with permission of Dr. Hans Joenje, VU Medical Center, Amsterdam, Netherlands | DMEM with 4.5 g/dL glucose, 10% FBS, penicillin (100 units/mL), streptomycin (100 mg/mL) |
| SCC4 | DSMZ-German Collection of Microorganisms and Cell Cultures GmbH |  |
| UD-SCC-2 | Dr. Thomas Carey, with permission of Dr. Henning Bier, Technical University Munich, Munich, Germany |  |
| UPCI: SCC-090 | American Type Culture Collection |  |
| TU-138 | Dr. Jennifer Grandis, UCSF | DMEM/F12 (1:1) |
| HN30 | Dr. Ravi Salgia, City of Hope | DMEM, 10% FBS, penicillin (100 units/mL), streptomycin (100 mg/mL) and NEAA |
| UM-SCC-1  UM-SCC-6 | Millipore | EMEM medium supplemented with NEAA, 10% fetal bovine serum (FBS), penicillin (100 U/ml) and streptomycin (100 μg/ml) |
| SCC-1483 | Dr. Lawrence Marnett, Vanderbilt University | DMEM with 4.5 g/dL glucose, 10% FBS, 1% hydrocortisone, penicillin (100 units/mL), streptomycin (100 mg/mL) |
| UM-SCC-47  UM-SCC-104  UM-SCC-22B | Millipore |  |

Supplementary table 2: Oligonucleotides List

| shEPHA2#1 For | | caccGCGTATCTTCATTGAGCTCAAtcaagagTTGAGCTCAATGAAGATACGC |
| --- | --- | --- |
| shEPHA2#1 Rev | | aaaaGCGTATCTTCATTGAGCTCAActcttgaTTGAGCTCAATGAAGATACGC |
| shEPHA2#2 For | | caccTCGGACAGACATATAGGATATtcaagagATATCCTATATGTCTGTCCGA |
| shEPHA2#2 Rev | | aaaaTCGGACAGACATATAGGATATctcttgaATATCCTATATGTCTGTCCGA |
| shROBO1 #1 For | | caccGCAGAAATACAGTCACATTATtcaagagATAATGTGACTGTATTTCTGC |
| shROBO1#1 Rev | | aaaaGCAGAAATACAGTCACATTATctcttgaATAATGTGACTGTATTTCTGC |
| shROBO1 #2 For | | caccTGACACATGACGCCAGATAAAtcaagagTTTATCTGGCGTCATGTGTCA |
| shROBO1 #2 Rev | | aaaaTGACACATGACGCCAGATAAActcttgaTTTATCTGGCGTCATGTGTCA |
| EPHA2 coding For | | aaaaactcgagATGGAGCTCCAGGCAGCCCGC |
| EPHA2 coding Rev | | aaaaaggtaccGATGGGGATCCCCACAGTGTTCACC |
| ROBO1 coding For | | ttttagatctATGATTGCGGAGCCCGCTCACTT |
| ROBO1 coding Rev | | ttttcccgggcctgctgctgcGCTTTCAGTTTCCTCTAATTCTTCATTAT |
| ROBO1 FLAG Rev | | ctagagtcgcggccgctTCACTTGTCGTCATCGTCTTTGTAGTCtgctgctgcGCTTTCAGTTTCC |
| EPHA2 HA Rev | | atgatctagagtcgcggccgcTCAAGCGTAATCTGGAACATCGTATGGGTAtgctgctgcGATGGGGATCCCCACAGTG |
| EPHA2 K645R | | CCGGTGGCCATCAGGACGCTGAAAGCCGG |
| ROBO1 Y932F | | GAAGAGAAACGGACTTactagtACCttcGCGGGTATCAGAAAAGTAAC |
| ROBO1 Y1073F | | ATCAGGGCAGCCTACTCCTttcGCCACCACTCAGCTCATC |
| vab-1 RNAi For | | ttttggtaccGTTCTTGTTCCACGTGTCGTC |
| vab-1 RNAi Rev | | tttagatctCCACATTCCACAAGTACATCC |
| sax-3 RNAi For | | ttttggtaccTTCCGAAGTGAGTCTCTTCTC |
| sax-3 RNAi Rev | | tttagatctCACCACCAACAATCGAGCATG |

Supplementary table 3: Antibodies List

| Antibodies used |  |  |
| --- | --- | --- |
| Antigen | Company | Cat # |
| FLAG Clone M2 | sigma | F1804 |
| b-actin | sigma | A5441 |
| EPHA2 | Santa Cruz | SC924 |
| EPHA2 pS897 | Cellsignaling | 6347 |
| EPHA2 pY588 | Cellsignaling | 12577 |
| ROBO1 | Invitrogen | PA5-29917 |
| AKT pS473 | Cellsignaling | 9271 |
| EGFR pY1068 | Cellsignaling | 2236 |
| HA | Cellsignaling | 3724 |
